# Polygenic Scores for Plasticity: A New Tool for Studying Gene-Environment Interplay

**DOI:** 10.1101/2020.08.30.274530

**Authors:** Rebecca Johnson, Ramina Sotoudeh, Dalton Conley

## Abstract

Outcomes of interest to demographers—fertility; health; education—are the product of both an individual’s genetic makeup and his or her social environment. Yet Gene × Environment research (GxE) currently deploys a limited toolkit on the genetic side to study gene-environment interplay: polygenic scores (PGS, or what we call mPGS) that reflect the influence of genetics on levels of an outcome. The purpose of the present paper is to develop a genetic summary measure better suited for GxE research. We develop what we call *variance polygenic scores* (vPGS), or polygenic scores that reflect genetic contributions to plasticity in outcomes. The first part of the analysis uses the UK Biobank (N ~ 326,000 in the training set) and the Health and Retirement Study (HRS) to compare four approaches for constructing polygenic scores for plasticity. The results show that widely-used methods for discovering which genetic variants affect outcome variability fail to serve as distinctive new tools for GxE. Then, using the polygenic scores that do capture distinctive genetic contributions to plasticity, we analyze heterogeneous effects of a UK education reform on health and educational attainment. The results show the properties of a new tool useful for population scientists studying the interplay of nature and nurture and for population-based studies that are releasing polygenic scores to applied researchers.

## 1 Introduction

### 1.1 The growth of using genome-wide measures to study genetic moderation of environments

A wide range of research has shown how outcomes of interest to demographers—e..g, fertility; educational attainment; diseases with marked disparities such as obesity—are influenced by both an individual’s genetic makeup and his or her social environment. In turn, this research program, also called *gene × environment* (G×E) research, has undergone a large shift in how researchers summarize genetic variation.

Earlier research focused on how *single* or *small sets* of genetic variants moderated social environments to affect outcomes. These include studies of how polymorphisms in specific genes like *MAOA*, or the promoter region of 5-*HTTP*, moderate social conditions like stressful childhood experiences or parental abuse (e.g., Guo et al., 2008) (for a review, see Seabrook and Avison (2010)).

Two developments led researchers to abandon studying how small sets of genetic variants moderate environments. First was the failure of many single gene G×E studies to replicate (Duncan and Keller, 2011; Keller, 2014; Border et al., 2019). Second was growing evidence that most outcomes of interest to social and behavioral scientists—educational attainment; body mass index (BMI); depression—are “polygenic,” that is, the result of small contributions of many variants across the genome, rather than “monogenic”(Boyle et al., 2017). As a result, researchers have moved away from studying how single genes or small sets of genes moderate environments to using polygenic scores (PGS) that summarize genome-wide contributions. As Section 1.3 shows, PGS have become the workhorse tool that social scientists use when studying genetic moderation of environments. As a result, large social science cohort studies—the Health and Retirement Study (Ware et al., 2018); the National Longitudinal Study of Adolescent to Adult Health (Braudt and Harris, 2020; the Wisconsin Longitudinal Study; the Fragile Families and Child Wellbeing Study—have either already released or are considering releasing polygenic scores alongside their standard survey measures.

The proliferation of polygenic scores as the workhorse tool for studying genetic moderation of social environments raises the question: what genome-wide summary *should* researchers use? Until now, researchers have developed scores that are meant to predict the conditional mean of an outcome. One problem with then using these scores to study Gene by Environment interactions (G × E) is that the PGS used in the interaction term is constructed from a meta-analysis of levels effects across multiple cohorts that differ temporally and geographically. As a result, the PGS may be particularly ill-suited for GxE analysis since it is based on the extraction of a signal for a main effect that is common across the plausible range of environments with which researchers may seek to interact it (i.e. the multiple cohorts, countries and contexts on which it is based). By contrast, by estimating a vPGS based explicitly on variation as the estimand in the training, the score may capture signals of variation across environmental contexts. More broadly, the goal of the present paper is to expand social scientists’ methodological toolkit by presenting a new summary measure: scores summarizing genetic contributions to plasticity.

The remainder of the introduction proceeds as follows. First, we outline two distinct forms of genetic moderation of social environments (Section 1.2). The first is when the environment’s impact on some outcome depends on that individual’s *genetic propensity to attain that same outcome*–for instance, pre-K having a larger effect on academic outcomes among children with an already-high genetic propensity towards high educational achievement. Building on the discussion in (Domingue et al., 2020), we call this form of genetic moderation *moderation through dimming or amplifying*.

Second is when the environment’s impact on some outcome depends on that individual’s *propensity towards variability* in an outcome. We can call this form of moderation moderation through plasticity. We argue that the majority of GxE research implicitly uses the first model of genetic moderation (moderation via dimming or amplifying). By making these implicit choices explicit, we highlight that little existing research uses tools suited for measuring the second form of genetic moderation.

Next, we outline how variance polygenic scores (vPGS) may capture this second form of genetic moderation (Section 1.4). We show that researchers’ focus on using methods for detecting genetic contributions to plasticity have thus far largely used the methods to find “top hits,” or a limited set of single nucleotide polymorphisms (SNPs) significantly associated with variability. We show our paper fills a gap by using these methods to construct genome-wide summary measures useful for GxE research, complementing other recent calls for better methods to detect genetic moderation of social environments (Domingue et al., 2020) and applications of vPGS (Schmitz et al., 2021).

### 1.2 Implicit models of genetic moderation: outcome moderation versus variability

Past typologies of different types of gene-environment interactions have focused on differences in the *shape* of the interaction (e.g., Boardman et al., 2014; and Derringer et al. (2019). For instance, Boardman et al. (2014) and Derringer et al. (2019) each summarize three shapes of interactions: (1) diathesis-stress, where those with both a risky genotype and a highly-stressful environment have adverse outcomes; (2) vantage-sensitivity, where those with a less risky genotype and a low-stress environment have particularly good outcomes; and (3) cross-over or differential suspectibility, where those with a risky genotype have adverse outcomes in high-stress environments but also have some of the best outcomes in supportive environments (Boyce and Ellis, 2005; Ellis et al., 2011). Researchers investigating genetic moderation of environments distinguish between these shapes through both theory and the form the interaction effect takes—for instance, diathesis-stress having a crossover shape.

Yet shape is only one dimension of how genotypes can moderate environments. The second dimension, which occurs regardless of shape, is what form of genetic variation moderates the impact of the environment on some outcome. Here, we review two types.

#### 1.2.1 Moderation through dimming and amplifying

The first type of interaction, coined by Domingue et al. (2020), is moderation through an individual’s genotype dimming or amplifying an environment. This occurs when a social environment either impedes or removes an impediment to the expression of a genetically-influenced outcome. For instance, people may vary in their genetic propensity to complete formal schooling (Lee et al., 2018). However, in certain societies, there may be limited access to schooling for the population or some subgroup within the population (e.g., access to higher education was limited for women for much of the twentieth century in the U.S. and elsewhere). If that constraint is removed, individuals’ genetic propensities towards higher education that had enjoyed no avenue for expression can then become manifest. In this case, we would expect a significant interaction in a model where a person’s years of schooling is regressed on (1) an indicator for the cohorts impacted by education reform and (2) a summary measure of a person’s genetic propensity to complete formal schooling. The coefficient between the genetic summary measure and reform would be null or smaller in the pre-reform years; it would become significant and positive during the post-reform years.^1^

As another example, economic changes in the U.S. have removed caloric constraints for much of the population. Genetic predispositions towards higher BMI now interact with an altered food environment (Guo et al., 2015; Conley et al., 2016). Those with a genetic predisposition towards higher BMI, which stems from a genetic architecture in part related to regulation of appetite and impulse control and in part related to metabolism (Locke et al., 2015), are more highly impacted by the new food environment.

#### 1.2.2 Moderation through plasticity

The case of BMI, however, also highlights a different form of genetic variation that can interact with changes to the environment. Individuals may vary not only in their propensity towards *higher* or *lower* BMI, but also vary in their propensity towards *changes in BMI* in the face of environmental changes. Some individuals have genotypes that are *less buffering* of environmental changes. When the environment changes (in either direction), their BMI is likely to exhibit large changes. Other individuals have genotypes that are *more buffering* of environmental changes. When they enter a more calorie-rich or more calorie-restricted environment, their BMI is less likely to change because they adapt to that environment in ways that minimize changes, regardless of where they were on the BMI distribution at baseline. A genetic predisposition towards higher or lower levels of BMI might be very different than a genetic predisposition towards *changes* in BMI in the face of shifting environmental conditions.

We call this form of genetic moderation *moderation through plasticity*. Plasticity can take two forms. First is variation in *within-individual plasticity*, which is relevant for dynamic outcomes like BMI and depression that change as individuals progress through the life course. As we discuss in the Conclusion, estimating genetic contributions to *within-person* variability is complicated by the lack of data with both large-scale cohorst that have been genotyped and repeated measures across genotyped individuals. Second, and more immediately tractable, is *population-level variation in plasticity*. To make more concrete, consider a shock that affects BMIs in a population—for instance, neighborhood violence that leads to more sedentary activity. While one form of gene by environment interaction might predict that those with genetic propensities towards high BMI are most impacted by the change, a plasticity-focused interaction would instead find individuals, that regardless of their propensity towards higher or lower BMI, are most sensitive to the environmental shock.

### 1.3 Gene × environment research using genome-wide polygenic scores has largely focused on moderation using levels scores

The previous section showed that one form of genetic moderation of environments is *moderation through dimming or amplifying*: those with different propensities towards an outcome are differentially impacted by some environmental change. Yet in a particular context—changes to neighborhoods interacting with genotype to impact BMI; changes to education policy interacting with genotype to affect schooling—genetic predispositions towards greater variability may also play a role.

Yet researchers’ workhorse tool for studying genetic moderation of environments—polygenic scores for levels of an outcome—has inadvertently narrowed their focus to outcome moderation. Researchers use a three-step process when they develop and use these scores:

**Step one**: estimate separate linear regressions of some outcome (*Y*) in a large training sample, to develop weights that reflect each variant’s contribution to levels of that outcome

**Step two**: use the weights from step one to construct a polygenic score (PGS) in a separate sample

**Step three:** interact that polygenic score with some measure of “E” to study genetic moderation of environments

Researchers in step one have focused on genetic contributions to levels of an outcome, rather than genetic contributions to variability. Table 1, focusing on recent gene by environment studies that use polygenic scores, shows that the majority focus on how the impact of environments on some outcome vary among people with different propensities for that same outcome—e.g., the impact of neighborhood features on Type II diabetes having a larger impact on those with higher genetic propensities.

**Table 1:** What type of moderation do recent gene by environment studies examine? The table presents a (non-exhaustive) list of recent gene by environment studies that use polygenic scores as the measure of genotype. With the exception of the study in gray, all study outcome moderation, or how the impact of an environmental trigger or buffer on an outcome varies by an individual’s genetic propensity towards that same outcome.

| Study | Outcome | Environment | PGS used to examine moderation |
| --- | --- | --- | --- |
| Barcellos et al. (2018) | BMI | Education reform | BMI |
| Liu and Guo (2015) | BMI | Childhood and adult SES | BMI |
| Amin et al. (2017) | Educational attainment | Educational attainment | BMI |
| Trejo et al. (2018) | Educational attainment; job status | School SES; school stratification; environment-agnostic heterogeneity in school-level random slopes on PGS | Educational attainment |
| Schmitz and Conley (2017) | Educational attainment | Veteran status (instrumented with Vietnam draft lottery) | Educational attainment |
| Herd et al. (2019) | Educational attainment | Gender/cohort | Educational attainment |
| Robinette et al. (2019) | Type II diabetes | Neighborhood disorder | Type II diabetes |
| Domingue et al. (2017) | Depressive symptoms | Spousal loss | Subjective wellbeing; depression |
| Halldorsdottir et al. (2019) | Depression | Childhood abuse | Depression |
| Mullins et al. (2016) | Depression | Childhood stressful life events and trauma | Depression |
| Papageorge and Thom (2020) | Educational attainment | Family SES | Educational attainment |

With the exception of Domingue et al. (2017), who examine how a genetic risk score for wellbeing buffers the impact of the loss of a spouse on depression, nearly all studies examine the dimming or amplifying mechanism. Furthermore, this focus on one form of genetic moderation is often *implicit*, with the researchers stating that they are studying gene by environment interactions, rather than stated as an *explicit* estimand (Lundberg et al., 2020), with the researchers stating that they are studying a particular type of gene by environment interaction. The failure to make the specific type of moderation explicit has led to missed opportunities to examine other forms of moderation.

### 1.4 Variance polygenic scores as a tool for examining new forms of genetic moderation

The implicit focus on one form of genetic moderation of social environments stems from the reliance on one tool for G×E research: polygenic scores trained to predict levels of an outcome. We follow others’ recent calls to expand social scientists’ toolbox for studying genetic moderation of environments. Recently, Domingue et al. (2020) discuss “dimmer-type” gene-environment interactions, which corresponds with outcome moderation, and “lens-type” gene-environment interactions, which take a different form.^2^ They argue that while social scientists often frame GxE research as wanting to study lens-type interactions, social scientists’ reliance on polygenic scores for levels of an outcome might impede their progress. As they put it: “The selection of PGS effects for examining lens-type GxE may be particularly challenging in that we construct PGSs from GWASs that only include main effects of SNPs. If the environmental context of the participants in the GWAS sample used to construct the PGS is similar to that in the test sample used to estimate GxE then it is unlikely to include SNPs that demonstrate lens-type patterns as the main effects of these SNPs will be close to zero”(p. 10). This call suggests that better tools for either variability moderation or “lens-type” moderation are genome-wide summary measures (PGS) constructed from weights that more closely mirror theory behind GxE.

Here, we present one approach: constructing genome-wide summary measures from models that measure genetic contributions to variability in outcomes.^3^ In the language of statistical genetics, these models are called “vQTL analyses,” or models for detecting variance-affecting loci. In turn, researchers have developed a variety of approaches for detecting genetic contributions to variability. But thus far, the researchers have only used these approaches to find the “top SNPs”—a few SNPs that have the lowest p-values in regressions performed separately for each SNP. They have not yet used the weights from these models to construct genome-wide scores for plasticity.

Rönnegård and Valdar (2011) first coined the term vQTL to discuss genetic contributions to trait variability. One of the earliest attempts at vQTL analysis was Yang (2012), who operationalize variability as a person’s Squared Z-score of a trait—the person deviates from the mean of an outcome in either direction. Wang et al. (2019) and others use the classic Levene’s test, which examines whether the error variance significantly differs across subgroups—in the genetics case, across the three subgroups (AA, AB, and BB) at a given variant. Yet these attempts can lead to false positives when trying to distinguish between variants that affect the mean of an outcome and variants that affect the variance.

Two methods aim to control for this mean-variance conflation. (2018) use sibling pairs to examine how variation in the sibling pair’s combined count of minor alleles at a locus contributes to that sibling pair’s standard deviation in the trait, controlling for the sibling pair’s mean levels of a trait. (2018) decompose trait variance into two components–an “additive effect” and a “dispersion effect”—and argue that the latter provides a measure of “when a SNP has a variance effect beyond that which can be explained by a general mean-variance relationship”(p. 1613).^4^

There is a significant gap in the use of these methods to study gene-environment interplay. Researchers have used each method to find “top hit” loci that contribute to variability in traits like BMI (Yang, 2012; Conley et al., 2018; Young et al., 2018).^5^ Some such as Wang et al. (2019), Young et al. (2018), and have also interacted these highly significant single SNPs one by one with measures of social environments. No studies of which we are aware have explored whether the “variance weights” that these methods generate can be aggregated to produce what we call *variance polygenic scores*, or genome-wide summary measures of a person’s plasticity. vPGS can expand demographers’ toolbox for studying gene-environment interplay. Our study is the first to build and characterize the properties of this new tool.

### 1.5 Research goals/questions

1. What are best practices for building variance polygenic scores?
2. When we build these scores, do they reflect distinctive genetic contributions to variability in a trait, or are they too correlated with scores for levels of an outcome to serve as a new tool for gene-environment research?
3. Applying the scores to a real-world example (education reform in the UK), what forms of moderation do we see?

## 2 Methods

### 2.1 Estimating vPGS weights in the UK Biobank

To build the two types of scores for comparison—a typical PGS (hereafter: mPGS) for levels of an outcome; a vPGS for variability in an outcome—we use the UK Biobank, a dataset containing about 500,000 individuals from across the United Kingdom. The sample was limited to respondents who passed quality control and were of British ancestry, using information provided by the UK Biobank, leaving us with 408,219 in our final analytic sample. Further information regarding sample construction and quality control can be found in Online Supplement Section S.1.

This size of the UK Biobank allows us to divide the sample into training and test sets while still maintaining sufficient statistical power for fitting GWAS and vQTL. Training and test sets were produced by randomly sampling respondents. 80% of the British subsample of the UKB was included in the training set; the remaining 20% made up the test set.

We analyze four outcomes: height, body mass index (BMI), educational attainment, and number of children ever born, a measure of fertility. The inverse normal transformations of the outcomes were calculated. Traits were also z-scored to create a second set of dependent variables, used in the Squared Z-score analyses. Unless specified as z-scored, a trait/outcome should be assumed to be inverse normal transformed.

For each outcome, first a regression was run predicting the inverse normal of that outcome, such that the weights reflect the contribution of each genetic locus to the mean level of the outcome. These regressions were performed using the software PLINK (version 1.9), controlling for age, sex, array, and the first 40 PCs.^6^ We refer to these regression weights as weights for Levels PGS, and they correspond to the traditional tools used in G×E research.

A set of second identical regressions predict the Squared Z-score, rather than inverse normal, of the outcome, corresponding with the method for vQTL analysis discussed in (Yang, 2012). Again, age, sex, array, and the first 40 PCs were included as controls. Since the z-score is squared, values which are the same number of standard deviations above or below the mean will receive the same value. Thus, the regression predicts distance from the mean, rather than the mean-level itself, though, as we argue above, this will still be correlated with the mean. We refer to the weights and polygenic scores produced by these regressions as Squared Z.

Third, regressions were run for each outcome on the sibling subsample of the UK Biobank, which includes 19,294 white British sibling pairs, while controlling for the same set of covariates as above. For each sibling pair, the intra-sibling mean and SD were calculated. We then residualized the SD with the mean and used this new residualized standard deviation as our outcome variable. Since each sibling pair was represented twice in the data, we used only one member of each sibling pair in the final regressions. We refer to the weights and polygenic scores produced by this method as Sibling SD.

Fourth, a Mean-Variance vQTL analysis using Levene’s test for variance heterogeneity was run using OSCA (www.cnsgenomics.com/software/osca) (Wang et al., 2019).The Levene’s test does not estimate the effect size and standard error, but rather assesses the equality of variances between sample groups (in this case, those that do and do not have a given allele). Thus, following (Zhang et al., 2019), OSCA re-scales the test statistics (p value) to effect size and standard error using Z-statistics. We refer to the weights and polygenic scores produced by this method as as Levene’s. OSCA requires one to distinguish continuous from discrete controls. As such, we treated the binary variables sex and array as discrete and age and PCs as continuous.

Fifth, a Mean-Variance vQTL analysis using heteroskedastic linear mixed models was similarly used (Young et al., 2018). Their method produces additive (mean) and variance effects. It also allows us to derive what they term dispersion effects, which are variance effects that are independent of the mean effect. We use the weights produced by these dispersion effects in subsequent analyses, referring to them as HLMM. Age, sex, array, and the first 40 PCs were included as both mean and variance covariates.

Finally, to ensure that results comparing the different vPGS were due to true differences between the scores, and not due to differences that arise from the smaller sample size and lower precision in the sibling-based method, for every vQTL or GWAS analysis run on the full sample an analogous analysis was run on a randomly-chosen subsample, where the number of respondents was set to be equal to the number of sibling pairs in the UK Biobank.

### 2.2 Constructing vPGS in the Health and Retirement Study (HRS)

Using the weights from the previous step, we constructed vPGS in the Health and Retirement study. The HRS sample is restricted to (1) self-identified European Americans, who (2) pass the HRS preprocessing procedure and are within 2 standard deviations of the mean of the first two principal components of their racial/ethnic group. This leaves *N* = 10, 554 respondents in the genotyping sample. Then, we filter to respondents with at least one wave of BMI, a primary outcome, collected (`r*bmi`). *N* = 5, 744 respondents remained after this exclusion.^7^ The replication code contains details on the outcome variable construction; most notably, since the HRS is timevarying with several waves, we took the most recently observed value of the outcome for each respondent.

### 2.3 Analytic approach

#### 2.3.1 Relationship between mPGS/vPGS and levels of an outcome

We use three tools to explore whether vPGS can capture genetic contributions to variability in an outcome, distinct from genetic contributions to levels of an outcome.

First, we estimate the following linear regression, where *i* indexes a respondent, *PGS* indicates the levels PGS (mPGS) or a variance PGS (vPGS), and *Y* is levels of the outcome trait (converted to the standard normal scale). *X_i_* includes the first 5 principal components (PCs). Our coefficient of interest is *β*_1_—we expect the levels PGS to significantly predict levels of a trait. We also conduct a robustness check where, in addition to controlling for age and sex in the construction of the vPGS weights using the UKB, we also control for these covariates in the regression. We find no substantive differences with these additional covariates.

In turn, SNPs are a mix of four types: (1) SNPs that affect neither levels of an outcome nor variance in an outcome, (2) SNPs that affect levels of an outcome but not its variance, (3) SNPs that affect variance in an outcome but not levels of an outcome, and (4) SNPs that affect both variance in and levels of an outcome. For traits that are not normally distributed, isolating SNPs of the third type is made more difficult by the fact that any SNP that affects the mean of an outcome will also affect the outcome’s variance (Young et al., 2018). Here, we aim to construct plasticity scores that capture genetic contributions to variability, and that are therefore comprised of SNPs of type three (SNPs that affect variance but not the mean) purged of general mean-variance artifacts from non-normal distributions and SNPs of type four. Since the distribution of type three and type four SNPs should be constant across each of the vPGS we compare, we interpret a larger positive coefficient on the vPGS from a regression of levels of an outcome on the vPGS as evidence that the vQTL method is picking up either (1) a large share of type four SNPs relative to type three SNPs or (2) has weights that fail to adjust for the mechanical relationship between mean and variance. For these regressions, our samples are the HRS and the held-out test set of the UKB.

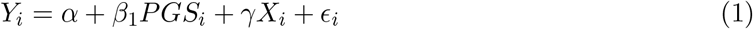

Second, we examine whether these patterns of correlation *after* constructing the vPGS in each sample are also present in the *underlying weights* that summarize each SNP’s contribution. We use linkage disequilibrium score (LD) score regression (Bulik-Sullivan et al., 2015b) for two purposes. First, we use the technique to compare the heritability of levels of an outcome to the heritability of plasticity in that outcome measured using the four techniques discussed above (Squared Z; Levene’s test; HLMM; sibling SD). Then, we compare the underlying genetic correlations between (1) levels and variability for each outcome, (2) across outcomes in levels and variability (Bulik-Sullivan et al., 2015b) (for a social science application of genetic correlations, see: (Wedow et al., 2018)).

Finally, since the analyses of heritability show very low heritability for vPGS like HLMM and sibling SD that are less confounded with levels of an outcome, we conduct two validation exercises to investigate whether the vPGS are capturing some form of plasticity and that the two scores do not just represent random noise. Using the HRS, which, unlike the UKB, has repeated measurements of the phenotype over time, we explore the relationship between each vPGS and two forms of plasticity. The first form of plasticity is *within-person variability*, which we measure using two versions of the within-person standard deviation in BMI: a version using raw values of BMI and a version detrended using age-specific trends. The second form of plasticity is *unexplained population-level variability*, which we operationalize by regressing BMI on age, sex, and the first five PCS, and then using the squared residual from that regression as the outcome.

Together, these analyses show that two of the polygenic scores for plasticity—one constructed using the Squared Z-score of an outcome; the other constructed using Levene’s test for variance heterogeneity—fail to summarize genetic contributors unique to variability in an outcome. However, two of the polygenic scores for plasticity—one summarizing “dispersion” effects; the other constructed from sibling variation—capture more distinctive genetic contributions. We focus on these two tools for the application we discuss in the next section, but include results with all four scores in the Online Supplement.

#### 2.3.2 Comparing mPGS versus vPGS as moderators of a UK education reform

We use these preferred plasticity scores to study heterogeneous effects of a large-scale education reform initiated in 1972 in England, Scotland, and Wales that extended how long students were legally required to stay in school from 15 to 16 years old (Barcellos et al., 2018). Using Barcellos et al. (2018)’s regression discontinuity design, we evaluate the extent to which the two effective vPGS are able to detect different forms of genetic moderation of this educational shock than the standard levels polygenic scores. We examine two different cases using the same reform as the exogenous environmental context: one where, following Barcellos et al. (2018), we examine the moderating role that the genetic risk of obesity plays in the relationship between education on body size and another where we evaluate the influence of genetic plasticity on downstream educational outcomes. Both outcomes are affected by gene and environment interactions but differ in the kinds of GxE effects they exhibit: while the effect for body size is the result of outcome moderation, the latter results from plasticity.

More specifically, following Barcellos et al. (2018), we use 2SLS to first instrument whether someone stayed in school until 16 years of age (Educ16) with whether they were younger than 16 when the reform went into place (post reform), and were therefore legally required to stay the extra year. This residualized version of Educ16 is then interacted in the second step with BMI PGS to evaluate whether there is an interaction between educational attainment and genes as they affect health outcomes. We also report a reduced form regression, where PGS is directly interacted with the post reform variable.

The main health outcome is Body Size - a weighted combination of BMI, waist-to-hip ratio, and body fat percentage. In a separate NBER preprint, (2019) identified the point in the Body Size distribution at which they should have the most power to detect an effect. We use this same distributional threshold to create an Above Threshold version of Body Size, the results for which can be compared to the continuous version of the outcome.

In a second set of analyses, we examine the role of genes in the impact of an additional year of education on downstream educational outcomes. Here, rather than instrumenting educational attainment, we look specifically at whether the effect of one’s genotype on education outcomes differs depending on whether one was born before or after the reform. This is akin to the reduced form regressions reported for BMI. We follow the previous literature (Barcellos et al., 2018), which found no effect of the reform on the likelihood of attending college, and focus on the outcomes of those who left school at the age of 18 or younger. We examine the effect of the reform on four educational outcomes, previously used in the literature. First, we consider whether the respondent left school at age 16 or later, since despite the reform some students still opted out of attending college through age 16 (Left School 16 or later). Second, we consider whether respondents achieved any certifications as a result of their education (Certification). For the last two outcomes, we explore whether they achieved specific certifications: O-levels or CSE (which were equivalent and later replaced by the GCSE in 1988) and A-levels.

Controls, for both sets of models, include a quadratic term for the number of days that passed from when the respondent was born until the time of reform (to factor out any time trends), dummy variables for the month born, sex, age at time of assessment in days, age squared, dummy variables for country of birth, the first 15 PCs, mPGS, the interaction between those PCs and Educ 16 (or, in the reduced form, Post Reform), and the interaction between mPGS with Educ16. Triangular kernel weights were used to assign more weight to observations closer to the reform and time trends were allowed to vary before and after the reform (Barcellos et al., 2018). Because we will show that the Squared Z-score and Levene’s test plasticity scores fail to capture variability distinct from mean effects, we do not present them in the main text but instead present them in the Supplementary Materials (Section S).

## 3 Results

### 3.1 How do the plasticity scores relate to levels of a trait?

The first question that arises when using plasticity scores for GxE research is: is the plasticity score simply capturing genome-wide contributions to levels of an outcome, rather than capturing genome-wide contributions to variability in an outcome? If the plasticity score looks very similar to social scientists’ standard tool for GxE research, it is less useful as a new tool for capturing distinctive forms of genetic moderation. Figure 1 summarizes the results of Model 1, or whether the plasticity score significantly predicts levels of an outcome. The left hand side shows the results in the smaller sample size HRS; the right panel the results in the larger sample size UK Biobank test set. Each bar represents one of the four outcomes of interest to demographers: height; BMI; education; number of children ever born (NEB).

**Fig. 1:**
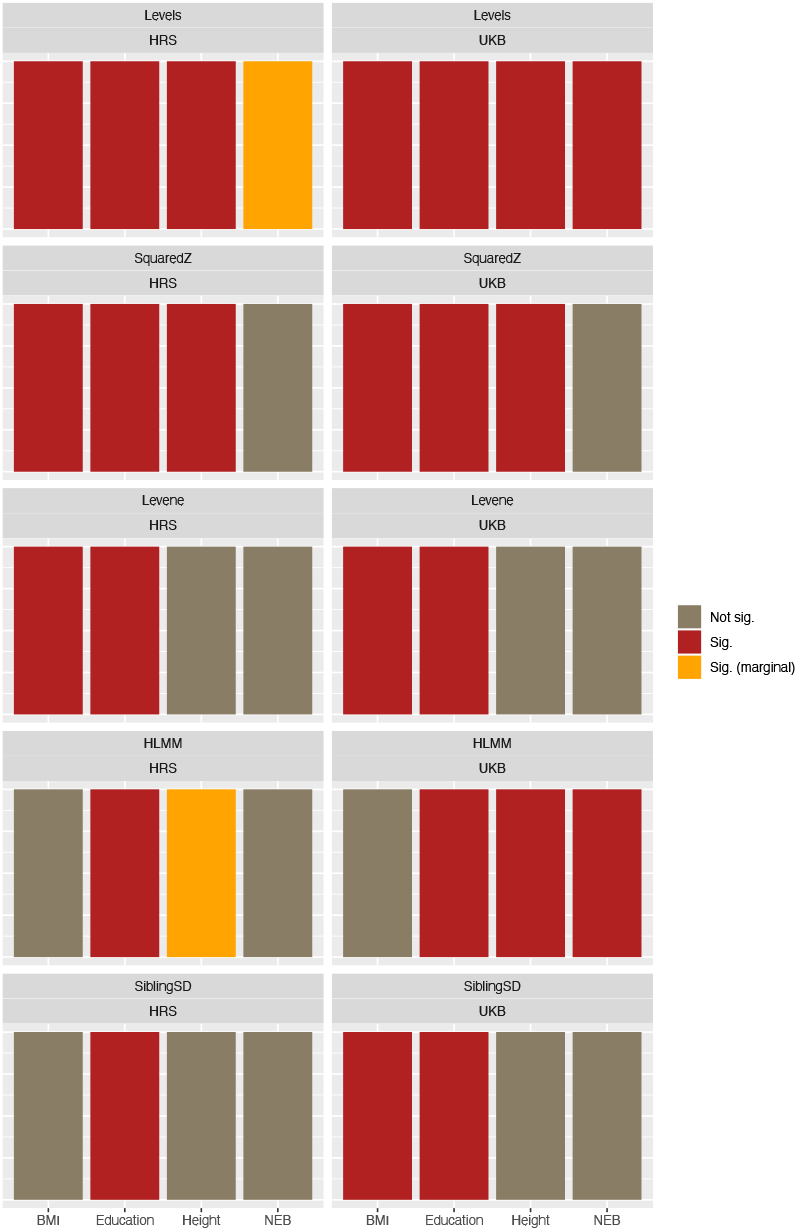
Significance of mPGS and plasticity scores in predicting levels of a trait. The figure shows results from the regression specified in Equation 1 in the HRS (left panel) and UKB test set (right panel). The top panel shows that, as expected, the levels PGS predict levels of an outcome (though the relationship with fertility in HRS is weaker than for height, BMI, and education). Moving downwards, across both samples, the Squared Z plasticity score performs the least well in that the plasticity score significantly predicts levels of an outcome for all outcomes except for number of children ever born. The Sibling SD score and HLMM perform best in the HRS test set

We see that, as expected, the levels PGS predict levels of an outcome. But in three out of the four traits, the Squared Z vPGS significantly predicts levels of a trait. In two of the four traits, the Levene’s test vPGS significantly predicts levels of a trait. In contrast, the sibling SD and HLMM vPGS were only significant for one out of the four traits in the HRS sample, though were significant for more traits in the UKB test set.

Overall, the results show that researchers hoping to use plasticity scores for gene-environment research should be careful to choose one of the tools that captures distinctive genetic contributions to plasticity apart from genetic contributions to an outcome’s mean. Online Supplement Section S.3 presents additional results, which include comparing the scores’ significance when we match the sample size of the non-sibling scores to the sample size in the sibling-based analyses.^8^

### 3.2 What is the genetic correlation between mPGS and vPGS?

The previous results show that when we aggregate weights from the different vQTL methods to produce a polygenic score for plasticity, some of the scores—most notably, the Squared Z vPGS— perform similarly to an mPGS in predicting levels of a trait. As a result, the score may be a less useful tool for examining certain forms of gene-environment interplay since they fail to capture distinctive genetic contributions to variability.

Here, we examine whether we can use tools aimed at using weights from mPGS to infer (1) heritability and (2) genetic correlation to examine the genetic architecture of plasticity.

Online Supplement Section S.5 contains the results from using LD score regression to examine the *univariate heritability* of each of the four outcomes—first, levels of an outcome (replicating previous work) and second, plasticity in that outcome (extending that work). We find that the only valid estimates of heritability are for the squared Z-score, possibly due to the method requiring weights with a certain degree of precision to generate non-zero heritability. Future research should investigate better methods for estimating heritability for less well powered vQTL weights.

Since the squared Z-score was the only one with non-zero heritability across outcomes, we examine the genetic correlation between (1) levels of each outcome (replicating past work by Bulik-Sullivan et al. (2015a)), (2) plasticity in each outcome (extending that work), and (3) levels and plasticity. Notably, these genetic correlations are prior to estimating the scores in a sample, so reflect a shared genetic architecture between contributors to levels of an outcome and contributors to variability in an outcome.

The top panel of Table 2 shows the genetic correlation between the levels PGS and the Squared Z vPGS for each of the outcomes. It shows that the weights for the levels PGS for that trait are significantly correlated with the weights for the Squared Z vPGS.

**Table 2:**
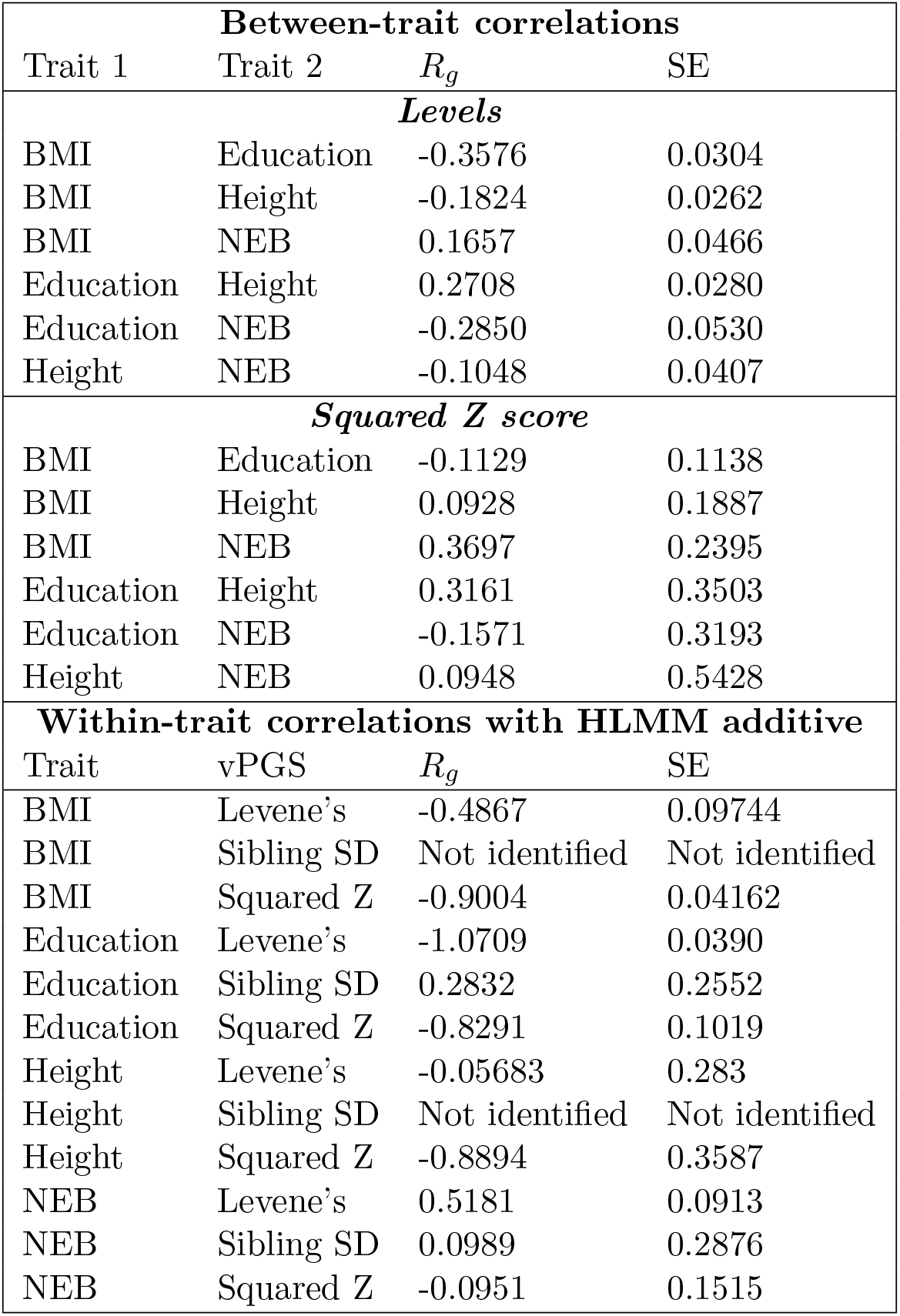
Genetic correlation results. The top panel shows the between-trait correlation in the standard levels weights. The middle panel shows the between-trait correlation in the squared Z vPGS weights, which are generally better powered. The bottom panel shows the within-trait correlation between: (1) the additive weights produced by the HLMM method, which are meant to identify mean effects purged of mean-variance correlations and (2) the non-HLMM vPGS.

| **Between-trait correlations** |  |  |  |
| --- | --- | --- | --- |
| Trait 1 | Trait 2 | $R_g$ | SE |
| ***Levels*** |  |  |  |
| BMI | Education | -0.3576 | 0.0304 |
| BMI | Height | -0.1824 | 0.0262 |
| BMI | NEB | 0.1657 | 0.0466 |
| Education | Height | 0.2708 | 0.0280 |
| Education | NEB | -0.2850 | 0.0530 |
| Height | NEB | -0.1048 | 0.0407 |
| ***Squared Z score*** |  |  |  |
| BMI | Education | -0.1129 | 0.1138 |
| BMI | Height | 0.0928 | 0.1887 |
| BMI | NEB | 0.3697 | 0.2395 |
| Education | Height | 0.3161 | 0.3503 |
| Education | NEB | -0.1571 | 0.3193 |
| Height | NEB | 0.0948 | 0.5428 |
| **Within-trait correlations with HLMM additive** |  |  |  |
| Trait | vPGS | $R_g$ | SE |
| BMI | Levene's | -0.4867 | 0.09744 |
| BMI | Sibling SD | Not identified | Not identified |
| BMI | Squared Z | -0.9004 | 0.04162 |
| Education | Levene's | -1.0709 | 0.0390 |
| Education | Sibling SD | 0.2832 | 0.2552 |
| Education | Squared Z | -0.8291 | 0.1019 |
| Height | Levene's | -0.05683 | 0.283 |
| Height | Sibling SD | Not identified | Not identified |
| Height | Squared Z | -0.8894 | 0.3587 |
| NEB | Levene's | 0.5181 | 0.0913 |
| NEB | Sibling SD | 0.0989 | 0.2876 |
| NEB | Squared Z | -0.0951 | 0.1515 |

**Table 3:** Impact of education reform on health outcomes

|  | HLMM vPGS |  | Sibling vPGS |  |
| --- | --- | --- | --- | --- |
|  | Body Size | Body Size (Threshold) | Body Size | Body Size (Threshold) |
|  | (1) | (2) | (3) | (4) |
| vPGS | −0.073<br>(0.049) | −0.044*<br>(0.018) | −0.018<br>(0.055) | −0.028<br>(0.021) |
| Educ16 (Instr.) | −0.225†<br>(0.133) | −0.110*<br>(0.050) | −0.231†<br>(0.134) | −0.111*<br>(0.051) |
| mPGS | 0.198***<br>(0.049) | 0.126***<br>(0.019) | 0.172**<br>(0.055) | 0.123***<br>(0.021) |
| vPGS x Educ16 (Instr.) | −0.017<br>(0.055) | 0.027<br>(0.021) | −0.024<br>(0.063) | 0.019<br>(0.024) |
| mPGS x Educ16 (Instr.) | 0.003<br>(0.056) | −0.087***<br>(0.021) | 0.017<br>(0.063) | −0.086***<br>(0.024) |
| Constant | −1.764***<br>(0.263) | −0.348***<br>(0.100) | −1.748***<br>(0.264) | −0.347***<br>(0.100) |
| Observations | 45,961 | 45,961 | 45,961 | 45,961 |
| R<sup>2</sup> | 0.061 | 0.026 | 0.054 | 0.023 |
| Adjusted R<sup>2</sup> | 0.061 | 0.025 | 0.053 | 0.022 |

*Note:*
\*p<0.1; \*\*p<0.05; \*\*\*p<0.01 †p < 0.1; \*p < 0.05; \*\* p < 0.01; \* \* \*p < 0.001

The middle panel of Table 2 shows *between-trait* patterns of genetic correlation for (1) the levels PGS^9^ and (2) the Squared Z vPGS. The analysis investigates whether there are similar patterns of cross-trait genetic correlation in variability in addition to levels. The results show that the patterns are similar except for the relationship between BMI and height. In particular, the one exception is that levels of height and BMI are negative genetically correlated (in other words, those with genetic propensities to be taller also have genetic propensities towards lower BMI, replicating the relationship in (2015a)) but plasticity in height and BMI is positively correlated. As we discuss in the Conclusion, this relationship deserves more attention in future research and could reflect differential sensitivity to environmental inputs to growth.

The bottom panel of the table also highlights the underlying genetic correlation between an mPGS meant to purge variance effects (the additive weights from the HLMM method) and each vPGS within a trait, which shows generally negative patterns. Appendix Section S.5 shows a visual summary of these correlations.

### 3.3 Validation that the vPGS correlates with plasticity

The previous sections showed that (1) especially the squared Z score vPGS was highly correlated with levels of an outcome and (2) that vPGS was the only one to have precise-enough estimates to be able to examine heritabilities and genetic correlations. Yet the noisiness of the vPGS estimates raises a question: could the results of Section 3.1 stem from scores that reflect random noise, rather than true contributions to variability?

Here, we report the results of the validation exercise discussed in Section 2.3.1. Figure 2 shows the results of relating each vPGS to within-person variability in BMI among respondents with at least three waves of BMI observations (Online Supplement Section S.6 discusses details of the sample construction and shows the full regression results). We see that individuals with higher vPGS have significantly more over-time variability in BMI than individuals with lower vPGS. Online Supplement Section S.6 also discusses a validation exercise where we regress the squared residual of BMI on each of the vPGS.

**Fig. 2:**
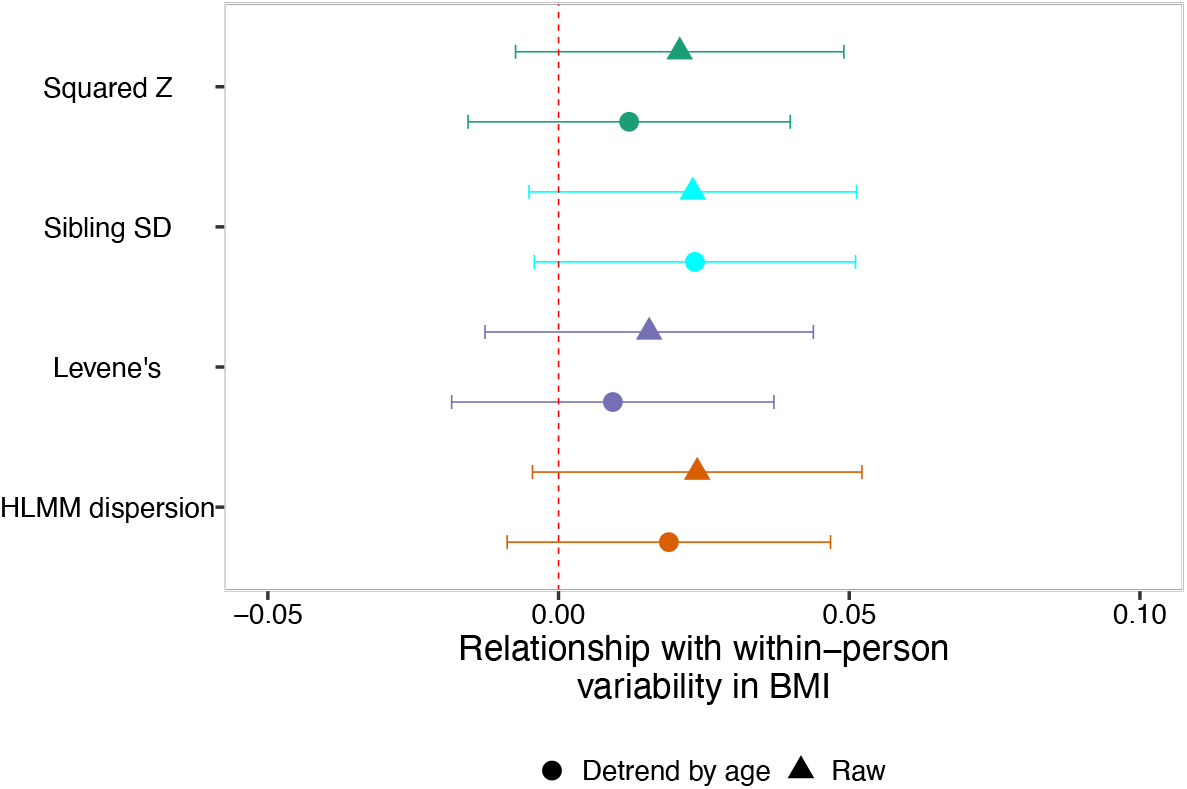
Relationship between vPGS and within-person variability in BMI: matched N sample. The figure, focusing on the sample where we restrict the estimation sample size for all scores to be equivalent to the estimation sample size for the sibling SD vPGS, shows two versions of the within-person variability analysis: a version with raw BMI over time and a version where BMI is detrended according to age patterns. The figure shows a general correlation between a respondent having a higher vPGS score and them having more variability in their BMI.

### 3.4 Summing up thus far: which vPGS can serve as new tools for GxE?

Taken together, the results show that the Squared Z vPGS vPGS is less useful for social scientists looking for a new tool to examine gene-environment interplay. The vPGS significantly predicts levels of an outcome across four diverse traits (height; BMI; education; number of children ever born). The Squared Z-score also exhibits patterns of underlying genetic correlation similar to those between levels of a trait. In contrast, the sibling standard deviation method (Conley et al., 2018) and dispersion weights (Young et al., 2018) show better properties in capturing distinctive genetic contributions to plasticity that appear less confounded with levels of an outcome.

Why might past research studying methods for vQTL have missed the ways in which certain methods fail to capture distinctive genetic effects? Section in the Online Supplement begins with the common way that researchers assess whether a method for detecting vQTLs overlaps with a method for detecting mean effects/normal QTLs: examining whether the two methods select similar SNPs as “top hits.”^10^ The results show that while the top hits comparison reveals some degree of overlap—for instance, the Squared Z score and Levene’s top hits display more overlap with the levels top hits than the other methods—this comparison might be too conservative. In particular, an mPGS and vPGS might not happen to have overlap in the SNPs with p values below a threshold but the two might have overlap in SNPs with non-zero weights that contribute to the final scores. Overall, the combined results show that social scientists interested in using vPGS as a new tool should look carefully at whether the vPGS is distinctive from, or nearly identical to, the mPGS for that outcome.

## 4 Using the vPGS to examine heterogeneous impacts of education reform

Having examined the properties of the different polygenic scores for plasticity, their distinctiveness from the standard tool for GxE research (mPGS), and their relationships to one another, we can use them to adjudicate between different mechanisms of genetic moderation. Specifically, we have argued that interactions between mPGS and the environment in predicting an outcome capture outcome moderation, while variance polygenic scores can capture a different form of heterogeneous effects.

With this in mind, we turn to using mPGS and different vPGSs in a practical example to explore which kind of genetic moderation is at play. We build upon the research of (2018). They investigate the impact of an educational reform that raised the required age of schooling from 15 to 16 years in England, Scotland, and Wales. Unlike measures like an individual’s own educational attainment or their parent’s educational attainment, which can lead to false-positive gene-environment interactions through confounding between the environmental shock and parent genotype (discussed in (Conley, 2016)), the reform’s timing is exogenous to genotype. It allows us to study the different forms of genetic moderation, as well as a chance to examine the performance of different vPGS measures in an applied example.

We evaluate two sets of outcomes. First, following (2018), we evaluate whether there was genetic moderation of the reform’s impact on health outcomes in the form of body size. If the form this moderation takes is *outcome moderation*, then the mPGS for BMI would significantly interact with the reform—the reform might have a larger impact on those with an already-low genetic propensity towards obesity (amplifying their advantage) or it might have a larger impact on those with a high genetic propensity (buffering their risk). Alternately, if the form this moderation takes is *variability moderation* (significant interaction between the vPGS and the post-reform indicator), the reform has larger impacts on those who, across many shocks, experience more swings in BMI.

Second, extending Barcellos et al. (2018), we evaluate whether there was genetic moderation of the reform’s impact on educational outcomes. Here, outcome moderation occurs if the reform has a larger impact on those with especially high or low genetic propensities towards educational attainment. Under the “education as the great equalizer hypothesis” (Barcellos et al., 2020), we might expect that those with the lowest educational polygenic score are the most impacted by the extra year of mandatory education. *Variability moderation* occurs if the reform has heterogeneous effects on individuals with different underlying genetic plasticity^11^.

### 4.1 Genetic moderation of the education reform’s impact on body size

For the body size models, presented in Table 4, the interaction between mPGS and being exposed to the reform is the only statistically significant result. None of the polygenic scores for plasticity show a significant interaction, with the exception of Squared Z-Score vPGS (Online Supplement Section S.8, which uncovers a marginally significant result (on the Above Threshold outcome)). This result is likely due to the high correlation between the Squared Z-Score vPGS and mPGS.

**Table 4:** Impact of education reform on health outcomes.

|  | Body Size<br>(1) | Body Size (Threshold)<br>(2) |
| --- | --- | --- |
| mPGS | 0.159***<br>(0.045) | 0.106***<br>(0.017) |
| Educ16 (Instr.) | -0.234<sup>†</sup><br>(0.134) | -0.112*<br>(0.051) |
| mPGS x Educ16 (Instr.) | 0.005<br>(0.051) | -0.074***<br>(0.019) |
| Constant | -1.728***<br>(0.264) | -0.342***<br>(0.100) |
| Observations | 45,961 | 45,961 |
| R<sup>2</sup> | 0.053 | 0.022 |
| Adjusted R<sup>2</sup> | 0.052 | 0.021 |

*Note:* <sup>†</sup> $p < 0.1$ ; \* $p < 0.05$ ; \*\* $p < 0.01$ ; \*\*\* $p < 0.001$

These results suggest that outcome moderation (rather than plasticity) is the main form that genetic moderation of the education reform takes when impacting these measures of health. Put differently, and as visualized in Figure 3, the reform has larger impacts on reducing obesity-related measures among those with already-higher genetic propensities towards obesity. The results largely replicate those found in Barcellos et al. (2018), and show that the mPGS the original authors used ended up corresponding to the type of genetic moderation that unfolded.

**Fig. 3:**
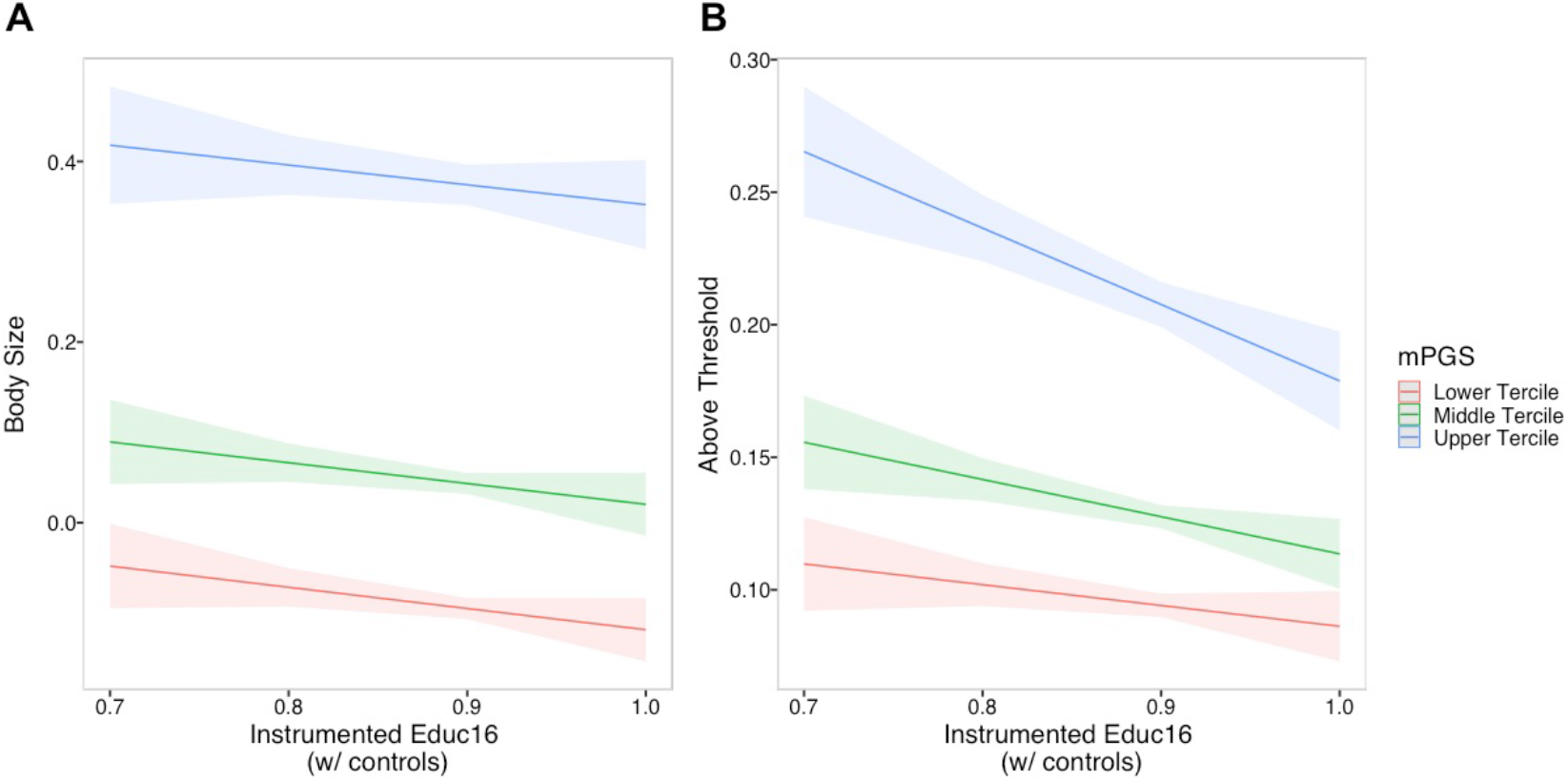
Interaction between mPGS and Instrumented Educ16 on Body Size outcomes.

### 4.2 Genetic moderation of the education reform’s impact on educational at-tainment

While the reform’s impact on health outcomes follows the pattern of outcome moderation, the reform’s impact on educational attainment might take a different form. When examining this impact, we find a significant interaction between the HLMM polygenic score for plasticity and Post Reform when predicting three of the four education outcomes: Left School 16 or later, Certification, and O-levels or CSE. The interactions between the HLMM plasticity score and Post Reform remain significant when controls are included. By contrast, we find significant interactions between mPGS and Post Reform for only one of the four outcomes: Left School 16 or later. The results are presented in Table 5 and visualized in Figure 4, which compares the predicted educational outcomes for children in the lowest, middle, and upper terciles of the HLMM vPGS distribution before and after the reform. For the significant interactions, the results show that those with higher HLMM polygenic score attain lower levels of education outcomes prior to the reform but equal levels of education outcomes after the reform, potentially because they had enhanced sensitivity to the positive effects of the reform. And because the results in Section 3.1 show that the HLMM plasticity scores capture genetic contributions to plasticity in outcomes distinct from genetic contributions to the conditional mean, we are more confident that the effect is a true positive.

**Table 5:** Genetic moderation of reform’s impact on educational attainment

| Controls? | *HLMM vPGS, Education Outcomes* |  |  |  |  |  |  |  |
| --- | --- | --- | --- | --- | --- | --- | --- | --- |
|  | Left School 16 or later |  | Certification |  | O-levels or CSE |  | A-levels |  |
|  | N | Y | N | Y | N | Y | N | Y |
|  | (1) | (2) | (3) | (4) | (5) | (6) | (7) | (8) |
| vPGS | −0.059***<br>(0.003) | −0.040***<br>(0.005) | −0.063***<br>(0.003) | −0.044***<br>(0.005) | −0.008<sup>†</sup><br>(0.004) | −0.007<br>(0.007) | −0.055***<br>(0.004) | −0.037***<br>(0.006) |
| Post Reform | 0.153***<br>(0.008) | 0.143***<br>(0.008) | 0.053***<br>(0.008) | 0.053***<br>(0.008) | 0.058***<br>(0.011) | 0.059***<br>(0.011) | −0.005<br>(0.009) | −0.005<br>(0.009) |
| mPGS |  | 0.022***<br>(0.005) |  | 0.021***<br>(0.005) |  | −0.0002<br>(0.007) |  | 0.021***<br>(0.006) |
| vPGS x Post Reform | 0.048***<br>(0.004) | 0.033***<br>(0.007) | 0.036***<br>(0.005) | 0.031***<br>(0.007) | 0.035***<br>(0.006) | 0.034***<br>(0.010) | 0.001<br>(0.005) | −0.003<br>(0.009) |
| mPGS x Post Reform |  | −0.017*<br>(0.007) |  | −0.005<br>(0.007) |  | −0.001<br>(0.010) |  | −0.003<br>(0.008) |
| Constant | 0.779***<br>(0.004) | 0.478**<br>(0.147) | 0.809***<br>(0.005) | 0.004<br>(0.156) | 0.557***<br>(0.006) | 0.247<br>(0.206) | 0.253***<br>(0.005) | −0.242<br>(0.178) |
| Observations | 25,690 | 25,690 | 26,012 | 26,012 | 26,012 | 26,012 | 26,012 | 26,012 |
| R<sup>2</sup> | 0.098 | 0.117 | 0.055 | 0.072 | 0.019 | 0.027 | 0.016 | 0.025 |
| Adjusted R<sup>2</sup> | 0.098 | 0.115 | 0.055 | 0.070 | 0.018 | 0.025 | 0.016 | 0.023 |

*Note:*
<sup>†</sup> $p < 0.1$ ; \* $p < 0.05$ ; \*\* $p < 0.01$ ; \*\*\* $p < 0.001$

**Fig. 4:**
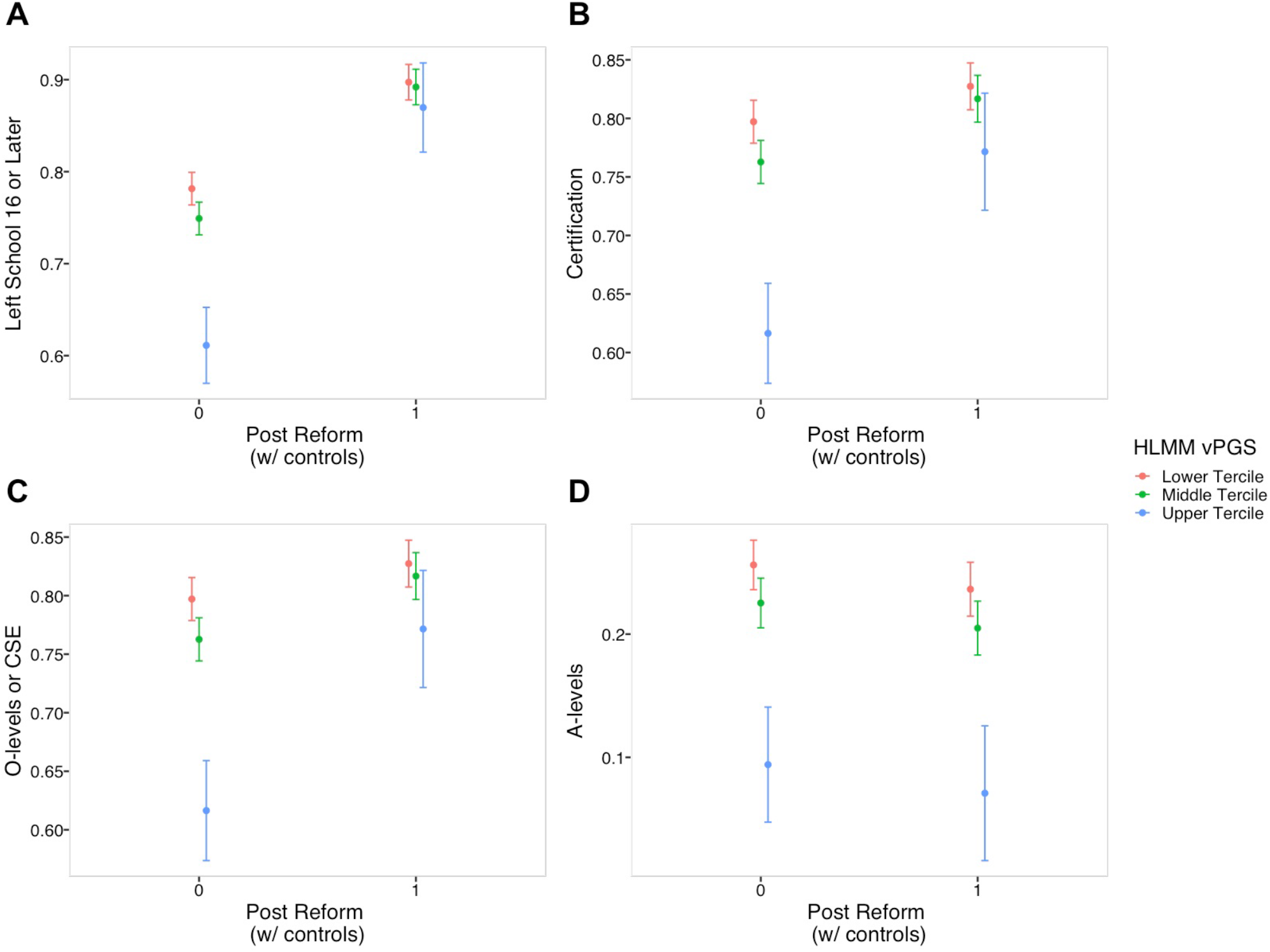
Interaction between HLMM polygenic score for plasticity and Post Reform on Educational Outcomes.

## 5 Discussion

Recognizing the biosocial nature of most outcomes of interest to demographers, social scientists are increasingly interested in how genetic variation moderates the impact of life-course events that range from society-wide education reforms to targeted policy interventions aimed at specific subgroups. That is, in addition to estimating direct main effects of genotypes and environments, social and behavioral scientists often seek to model the mutual dependence of nature and nurture. Numerous metaphors have been offered for this causal model of human traits—e.g., genetics as a lens (Domingue et al., 2020) or genetics as a prism refracting environmental influences into heterogeneous treatment effects (Conley and Fletcher, 2018).

In this paper, we argue that social scientists’ workhorse measure of *G* in GxE research—a genetic summary measure that reflects genetic contributions to levels of an outcome—commits those researchers to an implicit model of genetic moderation of environments. The model corresponds to what we call outcome moderation, and to what others have recently called “dimmer-type” moderation (Domingue et al., 2020). While this model may characterize some forms of gene-environment interplay, there are likely other forms of gene-environment interplay that summary measures constructed from aggregating effects on an outcome’s mean fail to capture.

We propose the use of polygenic scores for plasticity as an addition to social scientists’ toolbox. We first investigate the properties of this tool before applying it. First, focused on best practices, we show how conflation between genetic effects on an outcome’s mean and effects on that outcome’s variance begin with SNP-level analyses but then appear in the constructed scores. The conflation also makes it difficult to investigate whether plasticity in outcomes like BMI displays different patterns of heritability than levels of those outcomes, though an initial analysis of genetic correlations shows an interest flip where levels of BMI and height are negatively genetically correlated but plasticity in the two has a positive correlation. As a whole, we argue that researchers interested in a polygenic score for plasticity as a *distinctive* summary measure of genotype should be careful to construct scores based on weights from methods that try to adjust for false positive effects on the mean.

Second, applying the scores to a real-world application, we show how adding an *E* × *vPGS* analysis to an *E* × *mPGS* analysis can detect a particular type of GxE interaction that deploying only an mPGS would obscure. Building on (2018), we show that, in line with their results but contrary to our priors, *outcome moderation* best characterizes the education reform’s impact on health outcomes. But genetic plasticity might better explain the reform closing gaps in educational attainment between low and high-plasticity youth. These results show that one cannot know in advance with great certainty which form of moderation will be operative and thus researchers should test for both forms.

### 5.1 Limitations and directions for future research

The first limitation is that our application of the polygenic scores for plasticity was limited to one *E*: an education reform in the UK. In turn, and relevant to the theoretical discussion in Section 1.2, we might imagine that different environmental treatments are more or less likely to exhibit moderation by a person’s plasticity. Due to the issues others have raised about false positives where researchers think they are detecting GxE effects but instead are detecting unobserved confounding between genotype and environment (Conley, 2016; Domingue et al., 2020), we prioritized studying the effect of an “E” that was clearly causally identified over examining how multiple, potentially-confounded “E” interact with each of the focal vPGS. Future research leveraging other natural experiments that alter environments should incorporate plasticity scores to investigate their relevance for other contexts.

Second, as we outline in Section 1.2, there are at least two ways we can think about genetic contributions to plasticity. The first, *within-individual plasticity*, would require an estimation strategy where we train the inputs weights to the PGS on repeated measures of the same outcome within an individual (e.g., variation in BMI across many years). This form of plasticity is a promising avenue for future research. Predicting within-person (or within-family) variation over time without having to know explicitly what the fluctuating environmental factors are may prove to be a useful exploratory exercise before researchers try to hypothesize about specific factors in the environment that may be causing the fluctuation in genotypically-plastic individuals. Moreover, one could imagine using a within-person variability score to identify individuals who might be responsive to an intervention in advance—be that a drug trial or an educational intervention. The advantages of identifying such individuals include increased statistical power for the identification of effects in a pilot study before investing in a larger, more costly study. In terms of the feasibility of this second type of plasticity score, unfortunately, data sources like UKB that contain a large enough sample size to estimate new weights for polygenic scores lack large-scale repeated measures of the same individual. Once these data sources become available, future research can construct scores better designed for this form of plasticity.

Third, we might imagine two forms of plasticity. One form of plasticity is *trait specific* and occurs in response to various environmental triggers—so an individual with high BMI plasticity might have more BMI variability in response to many different environmental shocks (e.g., education reform; changes in food landscape; changes in peer group eating behavior). But another form of plasticity may be both *trait specific* and *environment specific*—so an individual may not have “generally high BMI plasticity,” but instead have high plasticity of BMI in response to a certain type of environmental trigger. The present approach to estimating variance-affecting SNPs weights and constructing *cross-environmental* vPGS captures the first type of plasticity, but fails to capture the second. For the second type of plasticity, researchers in statistical genetics are using flexible, machine learning methods to (1) focus on a specific “E” or environmental shock, (2) interact that “E” with many SNPs, (3) use regularization and other methods to find top-performing “SNP:E” interactions (for an early application noting challenges, see: (Boardman et al., 2014); more recently, (2016) use elastic net penalized regression to zero-out many of the SNP:“E” interactions). The weights from those methods focused on interactions between a specific “E” and each SNP could be used to estimate vPGS specific to certain environmental triggers.

Fourth, the present paper focuses still on scores that can be interacted with a specific measure of environment. But twin- and other pedigree-based approaches to estimating heritability have long been deployed to estimate GxE, for example, by assessing whether the additive heritability estimate changes in the face of differing social conditions (e.g., social class background; birth cohort)(e.g., Boardman et al., 2010; Vink and Boomsma, 2011). Newer methods such as GREML-based molecular methods allow for similar analysis under the same general framework where a shift in the additive (SNP) heritability is evidence of GxE (e.g., Rimfeld et al., 2018). However, since these methods each take an approach of variance decomposition, these methods cannot distinguish between different heritabilities due to a change in the genetic variance or a corresponding shift in the environmental variance. More importantly, by testing for changes in the total, additive heritability, as is the case for GxE studies using polygenic scores based on levels regressions, these approaches may be missing important GxE that do not result from differences in the predictive power of levels’ effects. One way to think about the plasticity or variance effect as it interacts with the environment is as an “environmental (and genetic)” epistasis term. That is, it is a non-additive effect that is not captured in traditional models. The goal of using vQTL methods is to capture this non-additive effect.

Finally, there is growing attention to how standard polygenic scores (mPGS) do not represent “pure” genetic measures of propensities; instead, the weights reflect a combination of direct genetic effects and biases from population stratification and genetic nurture (Kong et al., 2018; Trejo and Domingue, 2019; Zaidi and Mathieson, 2020). Because of these issues, for generating vQTL weights, the ideal design is either having the genotypes of two or more siblings along with the parent genotype, or having the genotype of three or more siblings and being able to add a fixed effect for the sibling pair. However, these methods require large-enough samples with one of those family-based structures. In the present paper, the sibling SD method provides one approach to addressing bias in the vQTL weights but future research should explore changes in vQTL weights when generated using a family-based design. In sum, the present article aims to equip demographers and social scientists with an additional tool for studying the interplay of genes and environment, one that captures a broader range of how these interactions play out in applied settings. As cohort studies make polygenic scores available to applied researchers, our paper suggests complementing the mPGS scores they are currently releasing with scores aimed at capturing genetic contributions to plasticity.

## Footnotes

1 Put differently, a polygenic score trained in societies where those constraints were attenuated or absent would poorly predict education before an expansion of schooling and then predict in an improved way — i.e. show increased genetic penetrance — once access to formal education was opened up.

2 As they describe: “When considering lenses, the relative effect of a given genotype may be positive for a ‘low’ level of the relevant environmental exposure and negative for ‘high’ levels of the exposure, or vice versa”(p. 9).

3 This approach complements the approach in Boardman et al. (2014) of studying genetic moderation of specific environmental shocks. In particular, in their study, they use what they call a genome-wide gene-by-environment interaction (GWGEI) approach that regresses level of an outcome (BMI) on each SNP’s interaction with an environmental moderator (education). They note the promise of the approach for capturing G×E, but also challenges with statistical power.

4 Other methods that we do not include in the present review because of their similarity to the four we focus on include the new deviation regression model (Marderstein et al., 2020), which models the absolute difference between an individual’s phenotype value and the phenotype medians within each genotype, and the double generalized linear model (DLGM) (**?**).

5 These are SNPs with effects on the outcome that fall below some p-value threshold.

6 Terms with interactions and higher order age variables (age^2^, age^2^ * sex), which have been employed in other studies, such as (Young et al., 2018), were excluded due to issues with multicollinearity.

7 We did this approach, rather than imputation, because the missingness was in the outcome variable rather than in a predictor.

8 This robustness check helps guard against us finding that the sibling SD score does not significantly predict levels of an outcome, while the non-sibling scores do, due to inadequate power for the sibling score compared to the scores estimated in a larger sample size. The fact that the patterns hold in the matched sample size supports our claim that the Squared Z and Levene’s test scores are less useful as distinctive tools.

9 These replicate results from Bulik-Sullivan et al. (2015a) for overlapping outcomes and also extend their analysis to look at additional outcomes like number ever born.

10 Researchers use this in conjunction with simulations comparing the methods, but those simulations likewise largely focus on one or two top causal SNPs.

11 Here, regressions that use vPGS also control for mPGS to ensure that the observed effects do not simply reflect outcome moderation. To understand how the inclusion of mPGS affects these results, we report regressions where mPGS is not controlled for in section S.8 of the SI. There, we also report the full regression results for the GxE analyses reported in the main text.

