## Supplementary Materials for "Polygenic Scores for Plasticity: A New Tool for Studying Gene-Environment Interplay"

### S Online Supplementary Materials

Here, we provide a guide to the online supplementary materials.

- Section S.1 discusses the score construction process in greater detail
- Section S.2 provides a descriptive summary of the bivariate correlation between: (1) the standard mPGS for each outcome, and (2) each of the four vPGS for each outcome, with Section S.2.1 focusing on HRS and Section S.2.2 on the UKB test set. The graphs show the strongest positive correlations for height and BMI, as well as the Squared Z and Levene's test vPGS.
- Section S.3 provides supplementary results for the regression of levels of an outcome on each mPGS and vPGS discussed in main text Section 3.1.
- Section S.5 discusses using LD score regression with the UKB vPGS weights to investigate the heritability of the different scores, discussing limitations of the method for detecting heritability in the scores with less confounding with levels of an outcome.
- Section S.6 discusses the results of the validation exercise to examine whether the vPGS are capturing any form of plasticity.
- Section S.7 provides supplementary results for the analysis of overlap in top hits discussed in main text Section 3.4.
- Section S.8 provides supplementary results for the GxE application discussed in main text Section 4.

#### S.1 vPGS construction process

##### S.1.1 Summary of raw count of SNPs for each score after estimation

**Table S1: Number of SNPS in raw vPGS estimation: UKB biobank** The main difference in $N$ SNPs is between (1) HLMM, which does pruning of SNPs as part of the estimation, and (2) methods that retain the majority of SNPs. Slight differences in the sample size across outcomes come from (1) the amount of missingness in the phenotype, which impacts the sample size (e.g., higher missingness in education) and (2) given that missingness among training set respondents, the missingness level in SNPs required to retain them in the estimation.

| score | outcome | n_snps |
|---|---|---|
| Levels | BMI | 729707 |
| Levels | Education | 729208 |
| Levels | Height | 729711 |
| Levels | NEB | 729705 |
| SquaredZ | BMI | 729707 |
| SquaredZ | Education | 729208 |
| SquaredZ | Height | 729711 |
| SquaredZ | NEB | 729705 |
| HLMM | BMI | 351996 |
| HLMM | Education | 351925 |
| HLMM | Height | 351988 |
| HLMM | NEB | 351974 |
| SiblingSD | BMI | 711962 |
| SiblingSD | Education | 709476 |
| SiblingSD | Height | 711983 |
| SiblingSD | NEB | 711928 |
| Levene | BMI | 777732 |
| Levene | Education | 776881 |
| Levene | Height | 777740 |
| Levene | NEB | 777731 |

##### S.1.2 UKB sample and HRS polygenic score construction from weights

The UK Biobank contains genotyping data for 488,377 respondents living in the United Kingdom between 2006 and 2010 (Bycroft et al. 2018). We restricted the sample to White British individuals using a designation provided as part of the UK Biobank data. White British individuals were those with self-reported race of white and an ethnicity of British. Finally, principal components of the genetic data were used to remove genetic outliers. The final sample of White British individuals includes 408,219 participants (see Bycroft et al. 2018 for more information on how White British measure was constructed).

Of the 488,377 participants, 438,427 were genotyped using the Applied Biosystems UK Biobank Axiom Array, which includes 825,927 markers, and the remaining 49,950 were genotyped using the UK BiLEVE Axiom Array by Affymetrix, which includes 807,411 markers. The arrays share 90% of the tagged genetic markers in common (Bycroft et al. 2018). The UK Biobank applied standard quality control (QC) measures to the genetic data in preparing it for dissemination and individuals who did not meet various QC analyses were labeled. Using their results, respondents putatively exhibiting sex chromosome aneuploidies, who had 10 or more relatives in the data or who were outliers with respect to missingness or heterozygosity were excluded from the final sample.

Information on the relatedness of respondents was provided by the UKB (Bycroft et al. 2018). The UKB inferred relatedness of participants using KING's robust estimator, a state-of-the-art algorithm for inferring relatedness from genetic data at scale that is robust to population structure (Manichaikul et al. 2010). The KING output, provided by the UKB, included kinship coefficients and identity-by-descent states, which were used to distinguish first degree relations (either sibling or parent-child) from 2nd and 3rd degree relations and full siblings from parent-child pairs, respectively, following thresholds provided Manachaikul et al. (2010). Of the 22,657 sibling pairs identified in total, 19,294 were White British and used in the sibling analyses.

As mentioned in the main text, the unrelated sample was divided randomly into training and test sets, composed of 80% and 20% of the sample, respectively. We then produced the genome-wide association measures (Levels, Sibling SD, Squared Z, Levene's, and HLMM).

After weights for these different measures were estimated, we used `plink` to construct variance polygenic scores in the test set of the UKB and the HRS. The UKB test set, like the training set, included only White British respondents. We filtered the HRS to European-American respondents and the sample excludes respondents with any relatedness.

While there are a variety of PGS construction methods, Ware et al. (2017) show how methods vary in three major ways: whether to use directly genotyped SNPs only or those and imputed SNPs; whether to "prune," or use a p-value threshold in the weights when estimating; whether and how to account for linkage disequilibrium (LD). Examining the impact of these choices in the HRS on aspects of polygenic scores like their $R^2$ values, the researchers show (1) slightly lower explanatory power when imputed SNPs are used rather than restricting to directly genotyped SNPs and (2) often better performance from not restricting to weights below a certain p-value threshold. Based on these results, and for simplicity in our method, we:

1. Use all weights rather than implementing a p-value threshold

2. Only use directly-genotyped SNPs

#### S.2 Relationship between mPGS and vPGS

##### S.2.1 Descriptive relationship between levels mPGS and vPGS: HRS

First, we look descriptively at the correlation between the levels PGS and each of the vPGS within HRS. Figure S1 shows the results for height, Figure S2 for BMI, Figure S3 for education, Figure S4 for number of children ever born.

**Fig. S1: Descriptive relationship between mPGS and vPGS: height (matched sample size)**

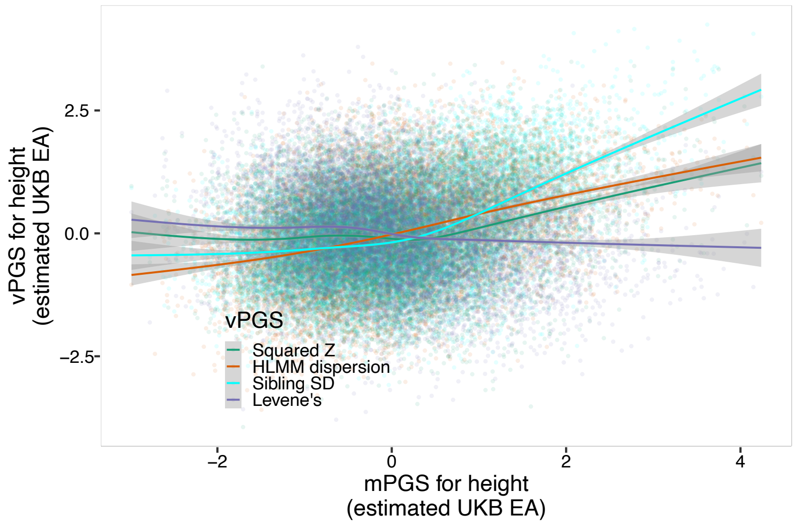


**Fig. S2: Descriptive relationship between mPGS and vPGS: BMI (matched sample size)**

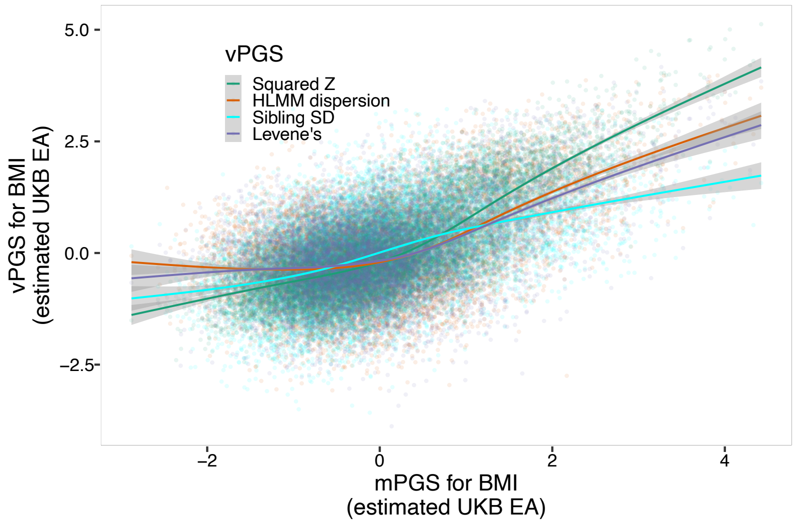

**Fig. S3: Descriptive relationship between mPGS and vPGS: education (matched sample size)**

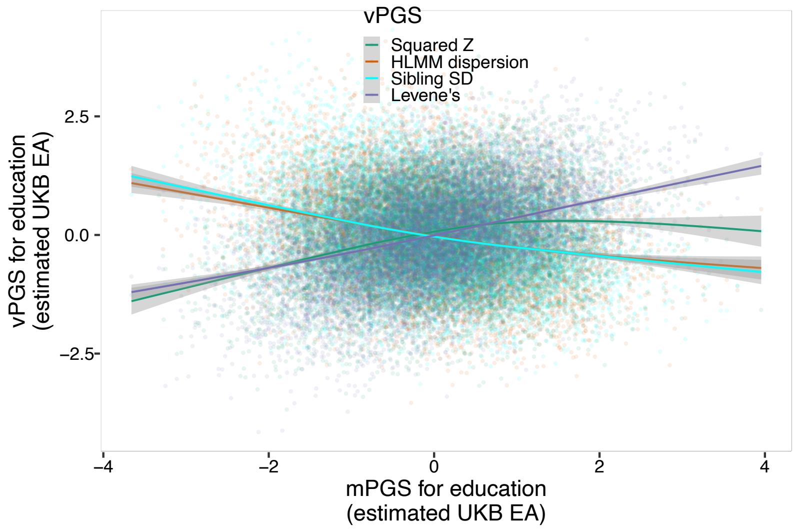


**Fig. S4: Descriptive relationship between mPGS and vPGS: number of children evern born (matched sample size)**

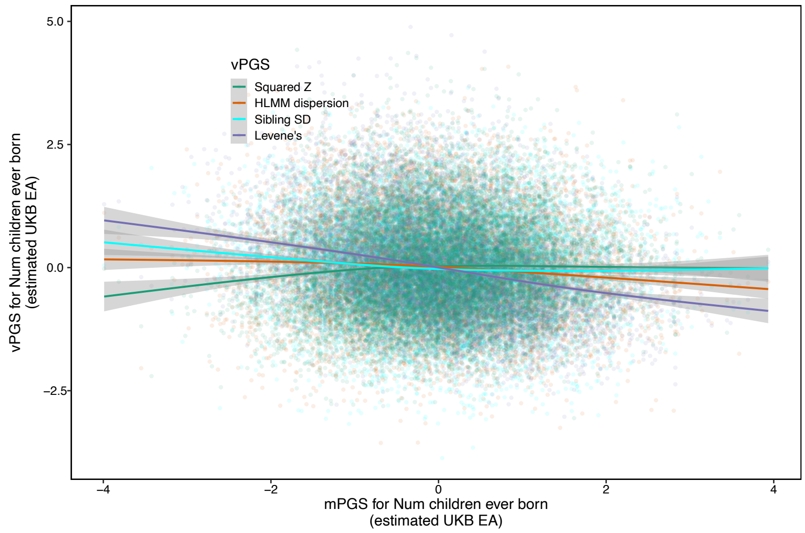

##### S.2.2 Descriptive relationship between levels mPGS and vPGS: UKB test set

Figure S5 - Figure S8 summarize the same descriptive relationships between the levels mPGS and the vPGS in the UKb test set.

**Fig. S5: Descriptive relationship between mPGS and vPGS: height (matched sample size)**

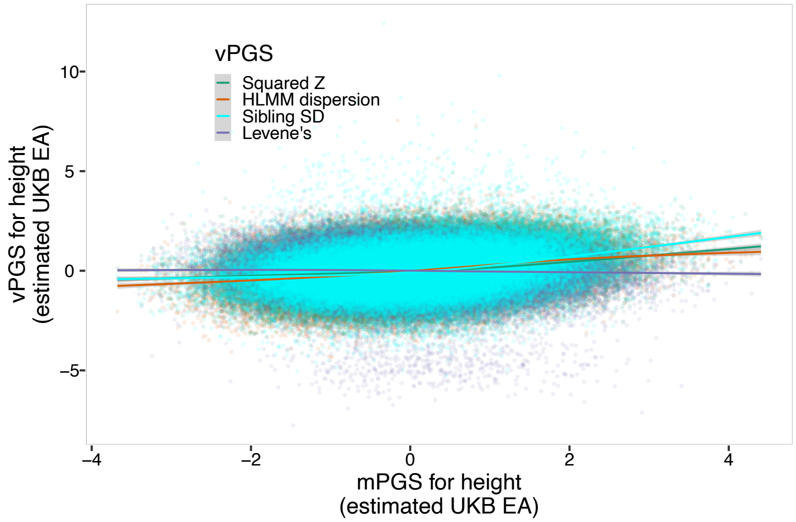


**Fig. S6: Descriptive relationship between mPGS and vPGS: BMI (matched sample size)**

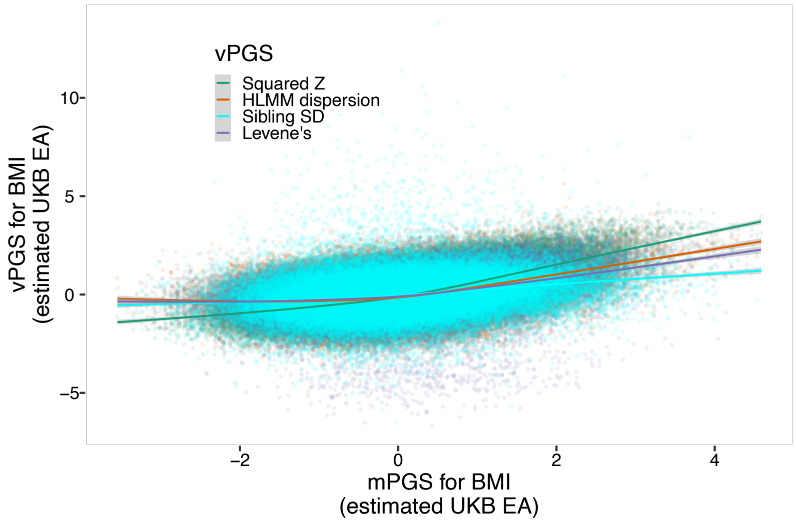

**Fig. S7: Descriptive relationship between mPGS and vPGS: education (matched sample size)**

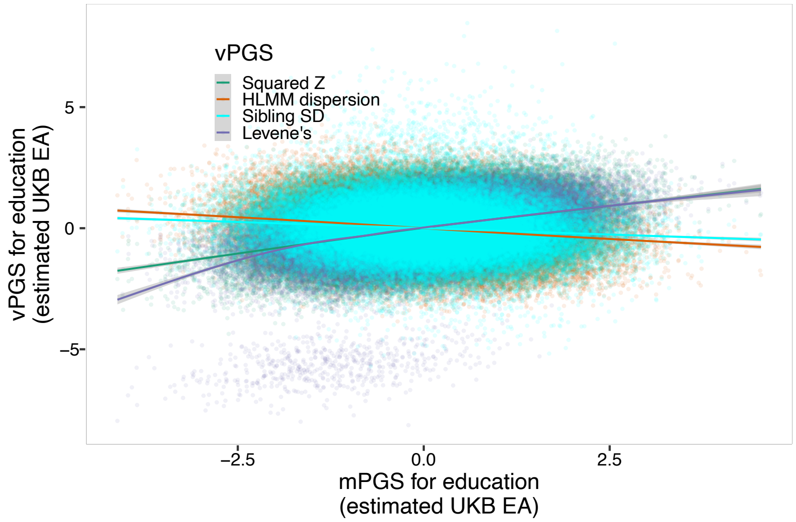


**Fig. S8: Descriptive relationship between mPGS and vPGS: number of children ever born (matched sample size)**

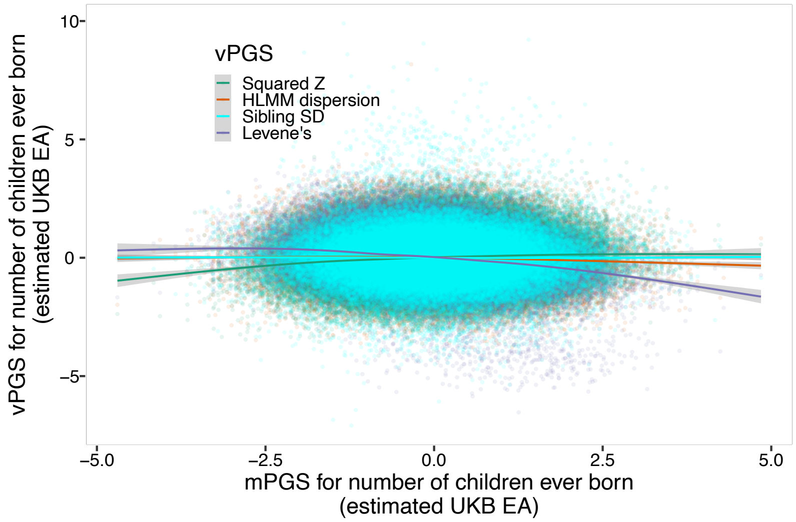


#### S.3 Relationship between vPGS and levels of an outcome

The Tables and figures that follow provide additional details of the results summarized in main text Section 3.1. First, Table S2 summarizes the significance of the regressions of levels on each of the mPGS/vPGS. The results show both those depicted in main text Figure 1 as well as the results from matching the sample size in the UKB from unrelated individuals to the sample size in the UKB used to estimate the sibling-based regression ($N \sim 22,000$). Section S.3.1 shows the coefficients from each set of scores in the HRS and UKB; it shows that scores based on weights estimated using a smaller sibling subsample are in the same direction but have wider confidence intervals (less precision) than scores based on weights estimated in the full sample. Finally, Section

S.4.1 presents the tables from the regressions.

**Table S2: Summarizing significance of relationship between PGS or vPGS and outcome in HRS and UKB test set** The table shows that Squared Z and Levene's have a higher rate of significantly predicting levels of an outcome than the HLMM and sibling SD vPGS. These results hold for when we (1) use the full sample size of unrelated individuals for the non-sibling vPGS, and (2) match the sample size for the non-sibling vPGS to the sample size for the sibling vPGS, which reduces statistical power/precision.

| | Levels | | Squared Z | | Levene's | | HLMM | | Sibling SD (same N for both) | |
|---|---|---|---|---|---|---|---|---|---|---|
| Phenotype | HRS | UKB (test set) | HRS | UKB (test set) | HRS | UKB (test set) | HRS | UKB (test set) | HRS | UKB (test |
| Height (matched N) | * (pos) | * (pos) | NS | * (pos) | NS | NS | * (pos) | * (pos) | NS | NS |
| Height (diff N) | * (pos) | * (pos) | * (pos) | * (pos) | NS | NS | NS (marginal and pos) | * (pos) | NS | NS |
| BMI (matched N) | * (pos) | * (pos) | NS | * (pos) | NS | * (neg) | NS | * (pos) | NS | * (pos) |
| BMI (diff N) | * (pos) | * (pos) | * (pos) | * (pos) | * (pos) | * (neg) | NS | NS | NS | * (pos) |
| Education (matched N) | * (pos) | * (pos) | NS | * (pos) | * (pos) | * (pos) | * (neg) | * (neg) | * (neg) | * (neg) |
| Education (diff N) | * (pos) | * (pos) | * (pos) | * (pos) | * (pos) | * (pos) | * (neg) | * (neg) | * (neg) | * (neg) |
| Number ever born (matched N) | NS (marginal and pos) | * (pos) | NS (marginal and pos) | NS | NS | NS | NS | NS | NS | NS |
| Number ever born (diff N) | NS (marginal and pos) | * (pos) | NS | NS | NS | NS | NS | * (pos) | NS | NS |

##### S.3.1 Regression results of levels of an outcome on each mPGS/vPGS (figures)

Across the figures, the solid lines represent the estimates when the UKB training sample size is allowed to vary across the sibling and non-sibling scores; the dashed lines represent the estimates when the sample size for the non-sibling scores is fixed to the sibling sample size.

**Fig. S9: Treatment effect of score on levels of outcome: height (HRS)**

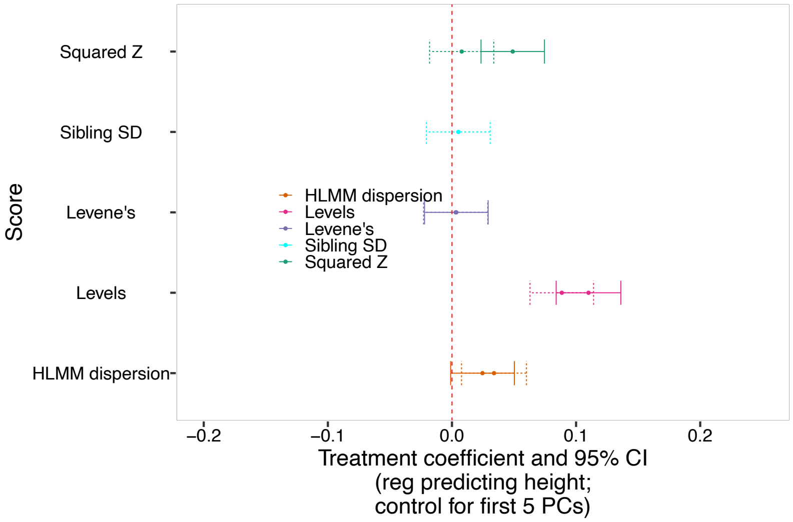


**Fig. S10: Treatment effect of score on levels of outcome: BMI (HRS)**

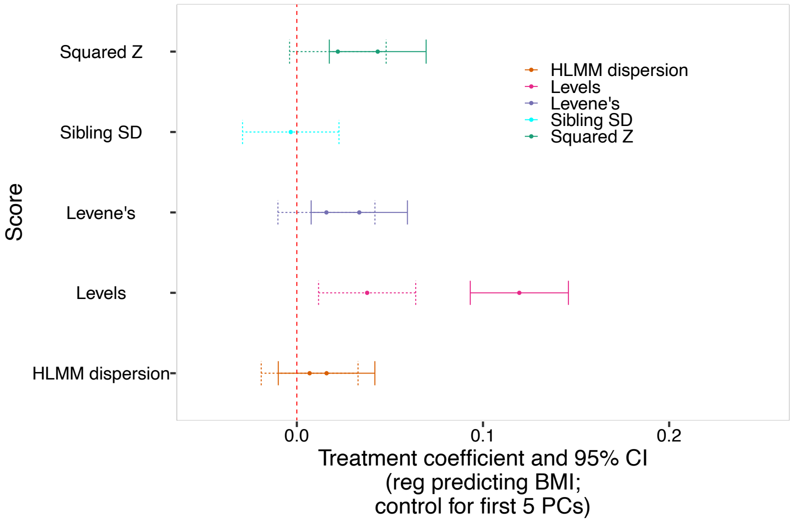

**Fig. S11: Treatment effect of score on levels of outcome: education (HRS)**

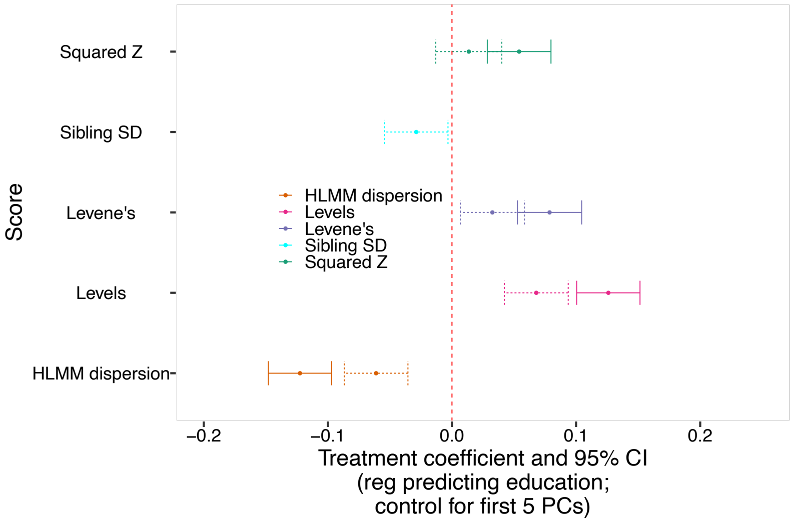


**Fig. S12: Treatment effect of score on levels of outcome: number of children ever born (HRS).**

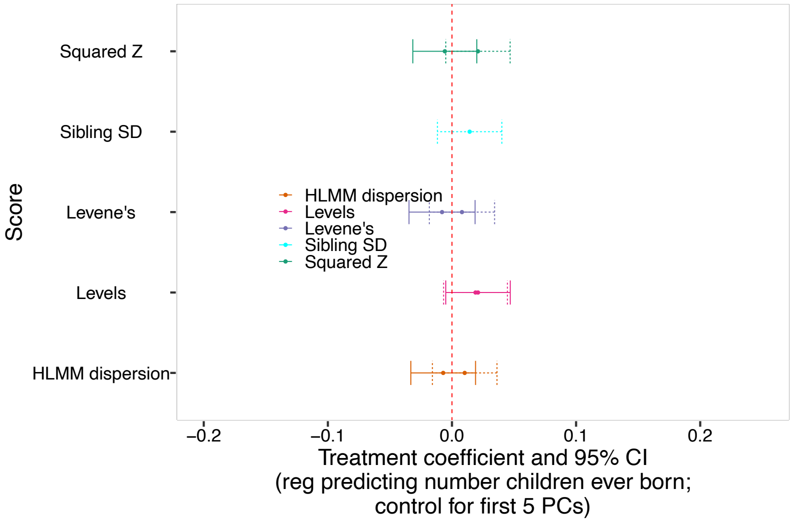

**Fig. S13: Treatment effect of score on levels of outcome: height (UKB)**

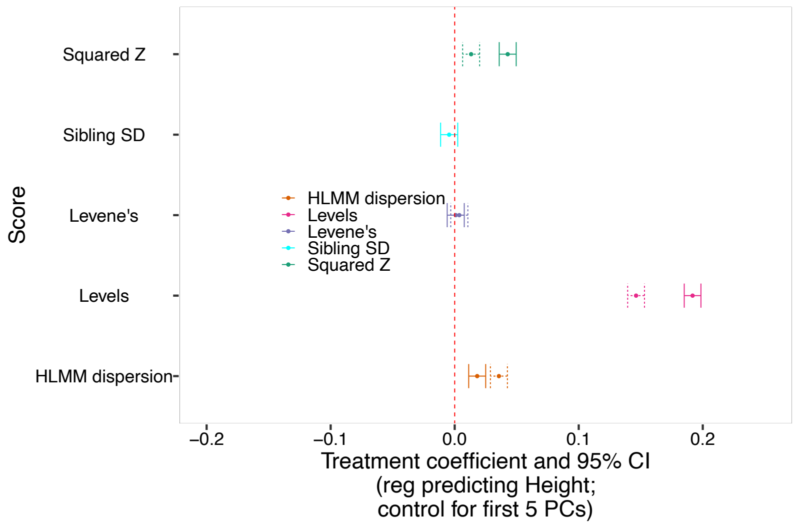


**Fig. S14: Treatment effect of score on levels of outcome: BMI (UKB)**

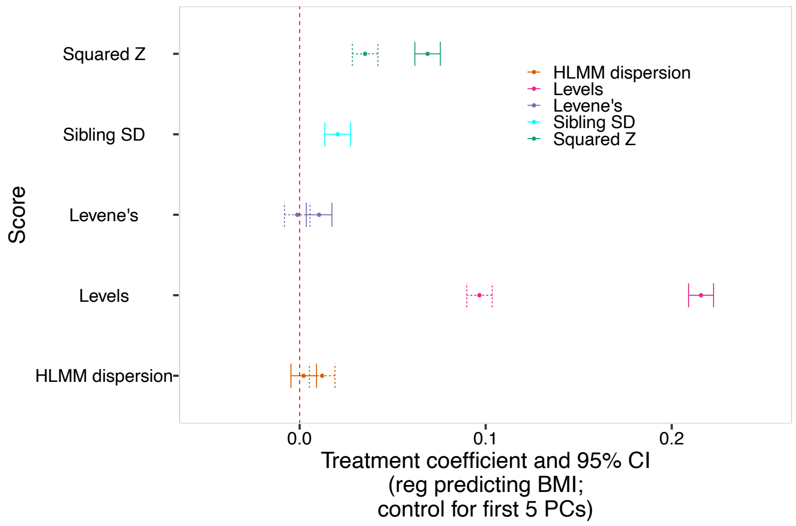

**Fig. S15: Treatment effect of score on levels of outcome: education (UKB)**

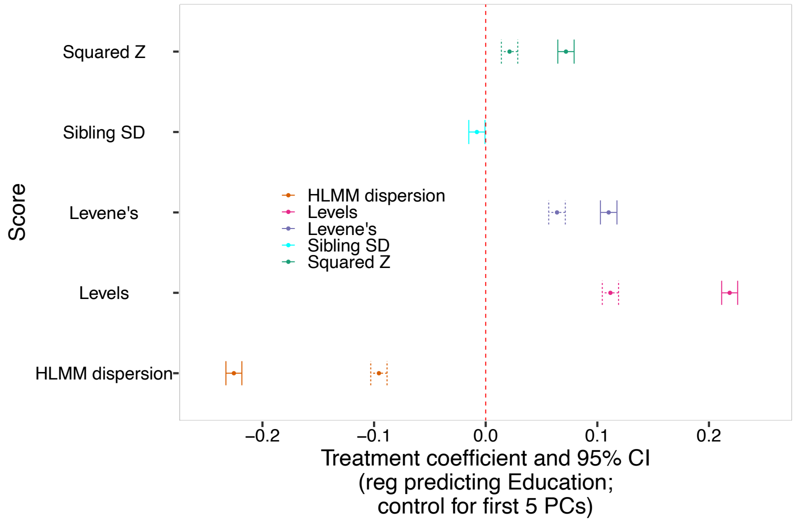


**Fig. S16: Treatment effect of score on levels of outcome: number of children ever born (UKB)**

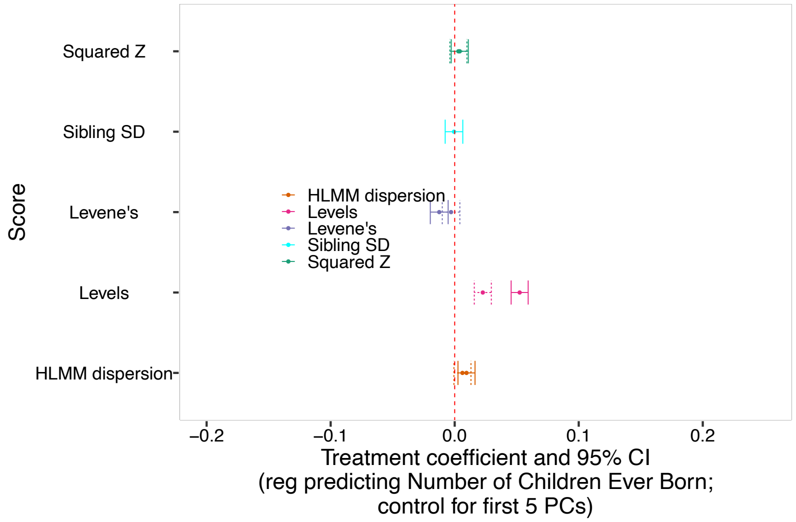

##### S.4 Robustness of vPGS scores

In their paper, Young et al. limit the set of SNPs for use in HLMM. Because we sought to replicate their procedure directly, we similarly limited the set of SNPs when running HLMM. However, this raises the potential issue that any differences between HLMM and the other methods may result from differences in the number of SNPs used when constructing scores. As such, we evaluated the robustness of the different methods to the number of SNPs used by correlating the scores that obtain when all SNPs are used with those that obtain when the Young et al. SNPs are used. The results can be found in Table S3.

**Table S3: Corr. between score with all SNPs vs. subset of SNPs used by Young et al.**

| Score | Outcome | Correlation |
|---|---|---:|
| Levels | BMI | 0.92 |
| Levels | Height | 0.97 |
| Levels | Education | 0.87 |
| Levels | NEB | 0.81 |
| SquaredZ | BMI | 0.96 |
| SquaredZ | Height | 0.82 |
| SquaredZ | Education | 0.76 |
| SquaredZ | NEB | 0.80 |
| SiblingSD | BMI | 0.88 |
| SiblingSD | Education | 0.90 |
| SiblingSD | Height | 0.88 |
| SiblingSD | NEB | 0.88 |
| Levene | BMI | 0.81 |
| Levene | Education | 0.47 |
| Levene | Height | 0.51 |
| Levene | NEB | 0.51 |

Though, in Yang (2012), traits were not transformed using the inverse normal transformation, we chose to use transformed traits to keep the trait distributions consistent across methods. To ensure the results are robust to this choice, in Table S4 we correlate, for each trait, the vGWAS and vPGS when traits are transformed (as used in the main text) with the vGWAS and vPGS when traits are not transformed. The results show a very high degree of correlation (greater than 0.9 in all instances).

**Table S4: Corr. between Levene's scores where traits are inverse normal transformed vs. not transformed**

| Trait | vGWAS-level Corr. | vPGS-level Corr. |
|---|---|---:|
| BMI | 0.93 | 0.96 |
| Height | 0.99 | 0.99 |
| Education | 0.97 | 0.97 |
| NEB | 0.95 | 0.96 |

###### S.4.1 Regression results of levels of an outcome on each mPGS/vPGS (tables)

**Table S5:** Regressing height on scores (HRS)

| score_combined | type | beta | se | t | p |
|---|---|---|---|---|---|
| Levels | Matched N | 0.09 | 0.01 | 6.77 | 0.00 |
| HLMM dispersion | Matched N | 0.03 | 0.01 | 2.54 | 0.01 |
| Sibling SD | Matched N | 0.01 | 0.01 | 0.38 | 0.70 |
| Squared Z | Matched N | 0.01 | 0.01 | 0.59 | 0.55 |
| Levene's | Matched N | 0.00 | 0.01 | 0.22 | 0.82 |
| Levels | Different N | 0.11 | 0.01 | 8.27 | 0.00 |
| HLMM dispersion | Different N | 0.02 | 0.01 | 1.88 | 0.06 |
| Levene's | Different N | 0.00 | 0.01 | 0.27 | 0.79 |
| Squared Z | Different N | 0.05 | 0.01 | 3.76 | 0.00 |

**Table S6:** Regressing levels of BMI on scores (HRS)

| score_combined | type | beta | se | t | p |
|---|---|---|---|---|---|
| Levels | Matched N | 0.04 | 0.01 | 2.84 | 0.00 |
| HLMM dispersion | Matched N | 0.01 | 0.01 | 0.52 | 0.60 |
| Sibling SD | Matched N | -0.00 | 0.01 | -0.25 | 0.80 |
| Squared Z | Matched N | 0.02 | 0.01 | 1.66 | 0.10 |
| Levene's | Matched N | 0.02 | 0.01 | 1.20 | 0.23 |
| Levels | Different N | 0.12 | 0.01 | 8.88 | 0.00 |
| HLMM dispersion | Different N | 0.02 | 0.01 | 1.21 | 0.23 |
| Levene's | Different N | 0.03 | 0.01 | 2.54 | 0.01 |
| Squared Z | Different N | 0.04 | 0.01 | 3.28 | 0.00 |

**Table S7:** Regressing levels of education on scores (HRS)

| score_combined | type | beta | se | t | p |
|---|---|---|---|---|---|
| Levels | Matched N | 0.07 | 0.01 | 5.17 | 0.00 |
| HLMM dispersion | Matched N | -0.06 | 0.01 | -4.67 | 0.00 |
| Sibling SD | Matched N | -0.03 | 0.01 | -2.21 | 0.03 |
| Squared Z | Matched N | 0.01 | 0.01 | 1.00 | 0.32 |
| Levene's | Matched N | 0.03 | 0.01 | 2.47 | 0.01 |
| Levels | Different N | 0.13 | 0.01 | 9.69 | 0.00 |
| HLMM dispersion | Different N | -0.12 | 0.01 | -9.41 | 0.00 |
| Levene's | Different N | 0.08 | 0.01 | 5.95 | 0.00 |
| Squared Z | Different N | 0.05 | 0.01 | 4.13 | 0.00 |

**Table S8:** Regressing number ever born on scores (HRS)

| score | beta | se | t | p | |
|---|---|---|---|---|---|
| Levels | Matched N | 0.02 | 0.01 | 1.45 | 0.15 |
| HLMM dispersion | Matched N | 0.01 | 0.01 | 0.77 | 0.44 |
| Sibling SD | Matched N | 0.01 | 0.01 | 1.08 | 0.28 |
| Squared Z | Matched N | 0.02 | 0.01 | 1.58 | 0.11 |
| Levene's | Matched N | 0.01 | 0.01 | 0.60 | 0.55 |
| Levels | Different N | 0.02 | 0.01 | 1.58 | 0.11 |
| HLMM dispersion | Different N | | -0.01 | 0.01 | -0.53 |
| 0.59 | | | | | |
| Levene's | Different N | -0.01 | 0.01 | -0.59 | 0.56 |
| Squared Z | Different N | | -0.01 | 0.01 | -0.44 |
| 0.66 | | | | | |

**Table S9:** Regressing BMI on scores (UKB)

| score_combined | type | beta | se | t | p |
|---|---|---|---|---|---|
| Levels | Matched N | 0.10 | 0.00 | 27.69 | 0.00 |
| HLMM dispersion | Matched N | 0.01 | 0.00 | 3.45 | 0.00 |
| Squared Z | Matched N | 0.04 | 0.00 | 10.04 | 0.00 |
| Levene's | Matched N | -0.00 | 0.00 | -0.37 | 0.71 |
| Sibling SD | Different N | 0.02 | 0.00 | 5.80 | 0.00 |
| Levels | Different N | 0.22 | 0.00 | 63.00 | 0.00 |
| HLMM dispersion | Different N | 0.00 | 0.00 | 0.62 | 0.54 |
| Squared Z | Different N | 0.07 | 0.00 | 19.66 | 0.00 |
| Levene's | Different N | 0.01 | 0.00 | 2.98 | 0.00 |

**Table S10:** Regressing educational attainment on scores (UKB)

| scores | type | beta | se | t | p |
|---|---|---|---|---|---|
| Levels | Matched N | 0.11 | 0.00 | 30.04 | 0.00 |
| HLMM dispersion | Matched N | -0.10 | 0.00 | -25.67 | 0.00 |
| Squared Z | Matched N | 0.02 | 0.00 | 5.70 | 0.00 |
| Levene's | Matched N | 0.06 | 0.00 | 16.84 | 0.00 |
| Sibling SD | Different N | -0.01 | 0.00 | -2.14 | 0.03 |
| Levels | Different N | 0.22 | 0.00 | 59.79 | 0.00 |
| HLMM dispersion | Different N | -0.23 | 0.00 | -61.91 | 0.00 |
| Squared Z | Different N | 0.07 | 0.00 | 19.24 | 0.00 |
| Levene's | Different N | 0.11 | 0.00 | 29.15 | 0.00 |

**Table S11:** Regressing height on scores (UKB)

| scores | type | beta | se | t | p |
|---|---|---|---|---|---|
| Levels | Matched N | 0.15 | 0.00 | 42.27 | 0.00 |
| HLMM dispersion | Matched N | 0.04 | 0.00 | 10.20 | 0.00 |
| Squared Z | Matched N | 0.01 | 0.00 | 3.79 | 0.00 |
| Levene's | Matched N | 0.00 | 0.00 | 1.06 | 0.29 |
| Sibling SD | Different N | -0.00 | 0.00 | -1.27 | 0.21 |
| Levels | Different N | 0.19 | 0.00 | 55.88 | 0.00 |
| HLMM dispersion | Different N | 0.02 | 0.00 | 5.19 | 0.00 |
| Squared Z | Different N | 0.04 | 0.00 | 12.24 | 0.00 |
| Levene's | Different N | 0.00 | 0.00 | 0.24 | 0.81 |

**Table S12:** Regressing number ever born on scores (UKB)

| scores | type | beta | se | t | p |
|---|---|---|---|---|---|
| Levels | Matched N | 0.02 | 0.00 | 6.46 | 0.00 |
| HLMM dispersion | Matched N | 0.01 | 0.00 | 1.78 | 0.07 |
| Squared Z | Matched N | 0.00 | 0.00 | 0.85 | 0.40 |
| Levene's | Matched N | -0.00 | 0.00 | -0.81 | 0.42 |
| Sibling SD | Different N | -0.00 | 0.00 | -0.16 | 0.88 |
| Levels | Different N | 0.05 | 0.00 | 14.95 | 0.00 |
| HLMM dispersion | Different N | 0.01 | 0.00 | 2.71 | 0.01 |
| Squared Z | Different N | 0.00 | 0.00 | 1.17 | 0.24 |
| Levene's | Different N | -0.01 | 0.00 | -3.40 | 0.00 |

**Table S13: Regressing number ever born on scores (HRS): matched N and including sex and age controls** The statistical significance does not differ from Table S8

| | *Dependent variable:* | | | | |
|---|---|---|---|---|---|
| | neb_stdnorm | | | | |
| | (1) | (2) | (3) | (4) | (5) |
| Levels mPGS (matched N) | 0.022<br>(0.013)<br>p = 0.079* | | | | |
| HLMM (matched N) | | 0.012<br>(0.013)<br>p = 0.348 | | | |
| Sibling SD (matched N) | | | 0.013<br>(0.013)<br>p = 0.315 | | |
| Levene's (matched N) | | | | 0.004<br>(0.013)<br>p = 0.774 | |
| Squared Z (matched N) | | | | | 0.015<br>(0.013)<br>p = 0.229 |
| PC1_5A | −2.030<br>(1.377)<br>p = 0.141 | −2.163<br>(1.377)<br>p = 0.117 | −2.171<br>(1.377)<br>p = 0.115 | −2.154<br>(1.379)<br>p = 0.119 | −2.151<br>(1.376)<br>p = 0.119 |
| PC1_5B | 2.539<br>(1.342)<br>p = 0.059* | 2.447<br>(1.342)<br>p = 0.069* | 2.432<br>(1.343)<br>p = 0.071* | 2.460<br>(1.346)<br>p = 0.068* | 2.411<br>(1.343)<br>p = 0.073* |
| PC1_5C | −0.213<br>(1.408)<br>p = 0.880 | −0.197<br>(1.409)<br>p = 0.889 | −0.055<br>(1.415)<br>p = 0.969 | −0.188<br>(1.409)<br>p = 0.894 | −0.211<br>(1.409)<br>p = 0.882 |
| PC1_5D | 0.520<br>(1.385)<br>p = 0.708 | 0.395<br>(1.383)<br>p = 0.776 | 0.512<br>(1.389)<br>p = 0.713 | 0.424<br>(1.389)<br>p = 0.761 | 0.432<br>(1.384)<br>p = 0.755 |
| PC1_5E | 6.172<br>(1.370)<br>p = 0.00001*** | 6.158<br>(1.373)<br>p = 0.00001*** | 6.186<br>(1.371)<br>p = 0.00001*** | 6.184<br>(1.383)<br>p = 0.00001*** | 6.219<br>(1.370)<br>p = 0.00001*** |
| Age | −0.017<br>(0.001)<br>p = 0.000*** | −0.017<br>(0.001)<br>p = 0.000*** | −0.017<br>(0.001)<br>p = 0.000*** | −0.017<br>(0.001)<br>p = 0.000*** | −0.017<br>(0.001)<br>p = 0.000*** |
| Female | | | | | |
| Constant | 33.274<br>(1.739)<br>p = 0.000*** | 33.243<br>(1.739)<br>p = 0.000*** | 33.222<br>(1.739)<br>p = 0.000*** | 33.222<br>(1.739)<br>p = 0.000*** | 33.184<br>(1.739)<br>p = 0.000*** |
| Observations | 5,743 | 5,743 | 5,743 | 5,743 | 5,743 |
| $R^2$ | 0.064 | 0.063 | 0.063 | 0.063 | 0.063 |
| Adjusted $R^2$ | 0.063 | 0.062 | 0.062 | 0.062 | 0.062 |
| Residual Std. Error (df = 5735) | 0.968 | 0.968 | 0.968 | 0.968 | 0.968 |
| F Statistic (df = 7; 5735) | 55.777*** | 55.439*** | 55.459*** | 55.317*** | 55.525*** |
| *Note:* | | | | | *p<0.1; **p<0.05; ***p<0.01 |

**Table S14: Regressing levels of height on scores (HRS): matched N and including sex and age controls** The statistical significance does not differ from Table S5

| | *Dependent variable:* | | | | |
|---|---|---|---|---|---|
| | height_stdnorm | | | | |
| | (1) | (2) | (3) | (4) | (5) |
| HLMM (matched N) | 0.030<br>(0.013)<br>p = 0.020** | | | | |
| Levels (matched N) | | 0.090<br>(0.013)<br>p = 0.000*** | | | |
| Sibling SD (matched N) | | | 0.008<br>(0.013)<br>p = 0.512 | | |
| Squared Z (matched N) | | | | 0.013<br>(0.013)<br>p = 0.318 | |
| Levene's (matched N) | | | | | 0.010<br>(0.013)<br>p = 0.428 |
| PC1_5A | 5.462<br>(1.358)<br>p = 0.0001*** | 5.224<br>(1.352)<br>p = 0.0002*** | 5.405<br>(1.358)<br>p = 0.0001*** | 5.419<br>(1.358)<br>p = 0.0001*** | 5.427<br>(1.358)<br>p = 0.0001*** |
| PC1_5B | −2.636<br>(1.325)<br>p = 0.047** | −2.903<br>(1.319)<br>p = 0.028** | −2.491<br>(1.324)<br>p = 0.060* | −2.449<br>(1.323)<br>p = 0.065* | −2.438<br>(1.324)<br>p = 0.066* |
| PC1_5C | 1.972<br>(1.395)<br>p = 0.158 | 1.598<br>(1.384)<br>p = 0.249 | 1.684<br>(1.390)<br>p = 0.226 | 1.710<br>(1.390)<br>p = 0.219 | 1.691<br>(1.390)<br>p = 0.224 |
| PC1_5D | 6.219<br>(1.365)<br>p = 0.00001*** | 6.318<br>(1.359)<br>p = 0.00001*** | 6.347<br>(1.366)<br>p = 0.00001*** | 6.354<br>(1.365)<br>p = 0.00001*** | 6.377<br>(1.367)<br>p = 0.00001*** |
| PC1_5E | 13.113<br>(1.368)<br>p = 0.000*** | 12.222<br>(1.360)<br>p = 0.000*** | 13.589<br>(1.352)<br>p = 0.000*** | 13.733<br>(1.358)<br>p = 0.000*** | 13.625<br>(1.352)<br>p = 0.000*** |
| rabyear | 0.018<br>(0.001)<br>p = 0.000*** | 0.018<br>(0.001)<br>p = 0.000*** | 0.018<br>(0.001)<br>p = 0.000*** | 0.018<br>(0.001)<br>p = 0.000*** | 0.018<br>(0.001)<br>p = 0.000*** |
| Female | | | | | |
| Constant | −34.944<br>(1.715)<br>p = 0.000*** | −35.096<br>(1.708)<br>p = 0.000*** | −35.015<br>(1.716)<br>p = 0.000*** | −35.033<br>(1.716)<br>p = 0.000*** | −35.038<br>(1.716)<br>p = 0.000*** |
| Observations | 5,744 | 5,744 | 5,744 | 5,744 | 5,744 |
| $R^2$ | 0.089 | 0.096 | 0.088 | 0.088 | 0.088 |
| Adjusted $R^2$ | 0.088 | 0.095 | 0.087 | 0.087 | 0.087 |
| Residual Std. Error (df = 5736) | 0.955 | 0.951 | 0.955 | 0.955 | 0.955 |
| F Statistic (df = 7; 5736) | 79.900*** | 87.107*** | 79.113*** | 79.202*** | 79.144*** |
| *Note:* | | | | | *p<0.1; **p<0.05; ***p<0.01 |

**Table S15: Regressing levels of BMI on scores (HRS): matched N and including sex and age controls** The statistical significance does not differ from Table S6

| | *Dependent variable:* | | | | |
|---|---|---|---|---|---|
| | bmi_stdnorm | | | | |
| | (1) | (2) | (3) | (4) | (5) |
| HLMM (matched N) | 0.013<br>(0.013)<br>p = 0.310 | | | | |
| Levels mPGS (matched N) | | 0.040<br>(0.013)<br>p = 0.002*** | | | |
| Sibling SD (matched N) | | | 0.005<br>(0.012)<br>p = 0.699 | | |
| Squared Z (matched N) | | | | 0.024<br>(0.013)<br>p = 0.054* | |
| Levene's (matched N) | | | | | 0.018<br>(0.013)<br>p = 0.142 |
| PC1_5A | −6.020<br>(1.341)<br>p = 0.00001*** | −6.091<br>(1.339)<br>p = 0.00001*** | −5.999<br>(1.341)<br>p = 0.00001*** | −6.027<br>(1.340)<br>p = 0.00001*** | −5.939<br>(1.340)<br>p = 0.00001*** |
| PC1_5B | −0.304<br>(1.306)<br>p = 0.816 | −0.531<br>(1.307)<br>p = 0.685 | −0.288<br>(1.306)<br>p = 0.826 | −0.380<br>(1.307)<br>p = 0.772 | −0.297<br>(1.306)<br>p = 0.821 |
| PC1_5C | −2.198<br>(1.372)<br>p = 0.110 | −2.117<br>(1.371)<br>p = 0.123 | −2.240<br>(1.372)<br>p = 0.103 | −2.200<br>(1.371)<br>p = 0.109 | −2.198<br>(1.371)<br>p = 0.110 |
| PC1_5D | 2.212<br>(1.347)<br>p = 0.101 | 2.148<br>(1.346)<br>p = 0.111 | 2.215<br>(1.349)<br>p = 0.101 | 2.196<br>(1.346)<br>p = 0.104 | 2.168<br>(1.347)<br>p = 0.108 |
| PC1_5E | −0.565<br>(1.336)<br>p = 0.673 | −0.862<br>(1.338)<br>p = 0.520 | −0.463<br>(1.335)<br>p = 0.729 | −0.695<br>(1.338)<br>p = 0.604 | −0.725<br>(1.344)<br>p = 0.590 |
| Age | 0.023<br>(0.001)<br>p = 0.000*** | 0.023<br>(0.001)<br>p = 0.000*** | 0.023<br>(0.001)<br>p = 0.000*** | 0.023<br>(0.001)<br>p = 0.000*** | 0.023<br>(0.001)<br>p = 0.000*** |
| Female | | | | | |
| Constant | −44.611<br>(1.693)<br>p = 0.000*** | −44.610<br>(1.692)<br>p = 0.000*** | −44.597<br>(1.694)<br>p = 0.000*** | −44.601<br>(1.693)<br>p = 0.000*** | −44.600<br>(1.693)<br>p = 0.000*** |
| Observations | 5,744 | 5,744 | 5,744 | 5,744 | 5,744 |
| $R^2$ | 0.112 | 0.113 | 0.112 | 0.113 | 0.112 |
| Adjusted $R^2$ | 0.111 | 0.112 | 0.111 | 0.111 | 0.111 |
| Residual Std. Error (df = 5736) | 0.943 | 0.942 | 0.943 | 0.943 | 0.943 |
| F Statistic (df = 7; 5736) | 103.457*** | 104.889*** | 103.315*** | 103.890*** | 103.639*** |
| *Note:* | | | | | *p<0.1; **p<0.05; ***p<0.01 |

**Table S16: Regressing levels of education on scores (HRS): matched N and including sex and age controls** The statistical significance does not differ from Table S7

| | *Dependent variable:* | | | | |
|---|---|---|---|---|---|
| | educ_stdnorm | | | | |
| | (1) | (2) | (3) | (4) | (5) |
| HLMM (matched N) | −0.059<br>(0.013)<br>p = 0.00001*** | | | | |
| Levels mPGS (matched N) | | 0.070<br>(0.013)<br>p = 0.00000*** | | | |
| Sibling SD (matched N) | | | −0.026<br>(0.013)<br>p = 0.042** | | |
| Squared Z (matched N) | | | | 0.012<br>(0.013)<br>p = 0.382 | |
| Levene's (matched N) | | | | | 0.035<br>(0.013)<br>p = 0.006*** |
| PC1_5A | 12.905<br>(1.375)<br>p = 0.000*** | 12.723<br>(1.373)<br>p = 0.000*** | 12.710<br>(1.377)<br>p = 0.000*** | 12.729<br>(1.378)<br>p = 0.000*** | 12.714<br>(1.376)<br>p = 0.000*** |
| PC1_5B | −1.605<br>(1.340)<br>p = 0.231 | −1.627<br>(1.339)<br>p = 0.225 | −1.678<br>(1.342)<br>p = 0.212 | −1.679<br>(1.345)<br>p = 0.213 | −1.706<br>(1.341)<br>p = 0.204 |
| PC1_5C | −0.773<br>(1.408)<br>p = 0.584 | −0.619<br>(1.406)<br>p = 0.660 | −0.635<br>(1.410)<br>p = 0.653 | −0.533<br>(1.409)<br>p = 0.706 | −0.660<br>(1.409)<br>p = 0.640 |
| PC1_5D | −1.268<br>(1.382)<br>p = 0.359 | −1.073<br>(1.380)<br>p = 0.437 | −1.343<br>(1.389)<br>p = 0.334 | −1.073<br>(1.384)<br>p = 0.439 | −0.979<br>(1.383)<br>p = 0.480 |
| PC1_5E | −5.261<br>(1.369)<br>p = 0.0002*** | −4.634<br>(1.369)<br>p = 0.001*** | −4.977<br>(1.370)<br>p = 0.0003*** | −4.799<br>(1.393)<br>p = 0.001*** | −5.218<br>(1.372)<br>p = 0.0002*** |
| Age | 0.015<br>(0.001)<br>p = 0.000*** | 0.015<br>(0.001)<br>p = 0.000*** | 0.015<br>(0.001)<br>p = 0.000*** | 0.015<br>(0.001)<br>p = 0.000*** | 0.015<br>(0.001)<br>p = 0.000*** |
| Female | | | | | |
| Constant | −29.186<br>(1.736)<br>p = 0.000*** | −29.340<br>(1.735)<br>p = 0.000*** | −29.223<br>(1.739)<br>p = 0.000*** | −29.256<br>(1.739)<br>p = 0.000*** | −29.333<br>(1.738)<br>p = 0.000*** |
| Observations | 5,744 | 5,744 | 5,744 | 5,744 | 5,744 |
| $R^2$ | 0.066 | 0.067 | 0.063 | 0.063 | 0.064 |
| Adjusted $R^2$ | 0.065 | 0.066 | 0.062 | 0.062 | 0.063 |
| Residual Std. Error (df = 5736) | 0.967 | 0.966 | 0.968 | 0.969 | 0.968 |
| F Statistic (df = 7; 5736) | 57.929*** | 59.181*** | 55.333*** | 54.818*** | 55.859*** |
| *Note:* | | | | | *p<0.1; **p<0.05; ***p<0.01 |

**Table S17: Regressing BMI on scores (UKB): matched N and including sex and age controls** The statistical significance does not differ from Table S9

| | *Dependent variable:* | | | | |
|---|---|---|---|---|---|
| | inv_norm_bmi | | | | |
| | (1) | (2) | (3) | (4) | (5) |
| HLMM (matched N) | −0.010***<br>p = 0.003 | | | | |
| Levels mPGS (matched N) | | 0.099***<br>p = 0.000 | | | |
| Sibling SD (matched N) | | | 0.010***<br>p = 0.004 | | |
| Squared Z (matched N) | | | | 0.036***<br>p = 0.000 | |
| Levene's (matched N) | | | | | 0.001<br>p = 0.747 |
| PC1 | −0.002<br>p = 0.524 | −0.002<br>p = 0.550 | −0.002<br>p = 0.496 | −0.002<br>p = 0.539 | −0.002<br>p = 0.515 |
| PC2 | −0.002<br>p = 0.511 | −0.003<br>p = 0.460 | −0.002<br>p = 0.515 | −0.002<br>p = 0.516 | −0.002<br>p = 0.511 |
| PC3 | −0.004<br>p = 0.292 | −0.003<br>p = 0.473 | −0.004<br>p = 0.286 | −0.004<br>p = 0.323 | −0.004<br>p = 0.291 |
| PC4 | 0.022***<br>p = 0.00001 | 0.019***<br>p = 0.00005 | 0.022***<br>p = 0.00001 | 0.021***<br>p = 0.00001 | 0.022***<br>p = 0.00001 |
| PC5 | 0.014***<br>p = 0.002 | 0.013***<br>p = 0.005 | 0.013***<br>p = 0.004 | 0.014***<br>p = 0.003 | 0.015***<br>p = 0.002 |
| Age | 0.008***<br>p = 0.000 | 0.008***<br>p = 0.000 | 0.008***<br>p = 0.000 | 0.008***<br>p = 0.000 | 0.008***<br>p = 0.000 |
| Sex | 0.242***<br>p = 0.000 | 0.242***<br>p = 0.000 | 0.242***<br>p = 0.000 | 0.242***<br>p = 0.000 | 0.242***<br>p = 0.000 |
| Constant | −0.803***<br>p = 0.000 | −0.809***<br>p = 0.000 | −0.804***<br>p = 0.000 | −0.804***<br>p = 0.000 | −0.803***<br>p = 0.000 |
| Observations | 81,400 | 81,400 | 81,400 | 81,400 | 81,400 |
| $R^2$ | 0.020 | 0.030 | 0.020 | 0.021 | 0.020 |
| Adjusted $R^2$ | 0.020 | 0.030 | 0.020 | 0.021 | 0.020 |
| Residual Std. Error (df = 81391) | 0.990 | 0.985 | 0.990 | 0.989 | 0.990 |
| F Statistic (df = 8; 81391) | 208.031*** | 310.654*** | 208.001*** | 220.583*** | 206.913*** |
| *Note:* | | | | | *p<0.1; **p<0.05; ***p<0.01 |

**Table S18: Regressing height on scores (UKB): matched N and including sex and age controls** The statistical significance does not differ from Table S11

| | *Dependent variable:* | | | | |
|---|---|---|---|---|---|
| | inv_norm_height | | | | |
| | (1) | (2) | (3) | (4) | (5) |
| HLMM (matched N) | 0.035***<br>p = 0.000 | | | | |
| Levels mPGS (matched N) | | 0.148***<br>p = 0.000 | | | |
| Sibling vPGS (matched N) | | | −0.009***<br>p = 0.0004 | | |
| Squared-z (matched N) | | | | 0.007***<br>p = 0.004 | |
| Levene's (matched N) | | | | | 0.006**<br>p = 0.023 |
| PC1 | 0.007***<br>p = 0.008 | 0.007***<br>p = 0.006 | 0.007***<br>p = 0.008 | 0.007***<br>p = 0.009 | 0.007***<br>p = 0.006 |
| PC2 | −0.004<br>p = 0.102 | −0.003<br>p = 0.227 | −0.004*<br>p = 0.096 | −0.004<br>p = 0.103 | −0.004<br>p = 0.103 |
| PC3 | 0.001<br>p = 0.769 | 0.002<br>p = 0.518 | 0.001<br>p = 0.810 | 0.001<br>p = 0.809 | 0.001<br>p = 0.807 |
| PC4 | −0.019***<br>p = 0.00000 | −0.019***<br>p = 0.000 | −0.018***<br>p = 0.00000 | −0.018***<br>p = 0.00000 | −0.018***<br>p = 0.00000 |
| PC5 | −0.050***<br>p = 0.000 | −0.052***<br>p = 0.000 | −0.049***<br>p = 0.000 | −0.050***<br>p = 0.000 | −0.050***<br>p = 0.000 |
| Age | −0.017***<br>p = 0.000 | −0.017***<br>p = 0.000 | −0.017***<br>p = 0.000 | −0.017***<br>p = 0.000 | −0.017***<br>p = 0.000 |
| Sex | 1.400***<br>p = 0.000 | 1.400***<br>p = 0.000 | 1.400***<br>p = 0.000 | 1.400***<br>p = 0.000 | 1.400***<br>p = 0.000 |
| Constant | −1.012***<br>p = 0.000 | −1.016***<br>p = 0.000 | −1.013***<br>p = 0.000 | −1.013***<br>p = 0.000 | −1.014***<br>p = 0.000 |
| Observations | 81,469 | 81,469 | 81,469 | 81,469 | 81,469 |
| $R^2$ | 0.508 | 0.529 | 0.507 | 0.507 | 0.507 |
| Adjusted $R^2$ | 0.508 | 0.529 | 0.507 | 0.507 | 0.507 |
| Residual Std. Error (df = 81460) | 0.700 | 0.685 | 0.700 | 0.700 | 0.700 |
| F Statistic (df = 8; 81460) | 10,514.760*** | 11,424.440*** | 10,466.870*** | 10,465.800*** | 10,465.010*** |
| *Note:* | | | | | *p<0.1; **p<0.05; ***p<0.01 |

**Table S19: Regressing levels of education on scores (UKB): matched N and including sex and age controls** The statistical significance does not differ from Table S10

| | *Dependent variable:* | | | | |
|---|---|---|---|---|---|
| | inv_norm_education | | | | |
| | (1) | (2) | (3) | (4) | (5) |
| HLMM (matched N) | −0.205*** | | | | |
| | p = 0.000 | | | | |
| Levels mPGS (matched N) | | 0.239*** | | | |
| | | p = 0.000 | | | |
| Sibling SD (matched N) | | | −0.022*** | | |
| | | | p = 0.008 | | |
| Squared Z (matched N) | | | | 0.048*** | |
| | | | | p = 0.000 | |
| Levene's (matched N) | | | | | 0.138*** |
| | | | | | p = 0.000 |
| PC1 | −0.033*** | −0.034*** | −0.035*** | −0.035*** | −0.027*** |
| | p = 0.0002 | p = 0.0001 | p = 0.0001 | p = 0.0001 | p = 0.002 |
| PC2 | −0.014* | −0.015* | −0.015* | −0.015* | −0.014* |
| | p = 0.087 | p = 0.068 | p = 0.082 | p = 0.077 | p = 0.099 |
| PC3 | 0.015* | 0.014* | 0.016* | 0.015* | 0.016* |
| | p = 0.077 | p = 0.092 | p = 0.066 | p = 0.072 | p = 0.064 |
| PC4 | −0.119*** | −0.116*** | −0.122*** | −0.121*** | −0.125*** |
| | p = 0.000 | p = 0.000 | p = 0.000 | p = 0.000 | p = 0.000 |
| PC5 | 0.070*** | 0.067*** | 0.072*** | 0.070*** | 0.056*** |
| | p = 0.000 | p = 0.000 | p = 0.000 | p = 0.000 | p = 0.00000 |
| Age | −0.040*** | −0.040*** | −0.040*** | −0.040*** | −0.040*** |
| | p = 0.000 | p = 0.000 | p = 0.000 | p = 0.000 | p = 0.000 |
| Sex | 0.220*** | 0.216*** | 0.219*** | 0.219*** | 0.222*** |
| | p = 0.000 | p = 0.000 | p = 0.000 | p = 0.000 | p = 0.000 |
| Constant | 3.421*** | 3.423*** | 3.412*** | 3.412*** | 3.407*** |
| | p = 0.000 | p = 0.000 | p = 0.000 | p = 0.000 | p = 0.000 |
| Observations | 71,195 | 71,195 | 71,195 | 71,195 | 71,195 |
| $R^2$ | 0.033 | 0.036 | 0.025 | 0.025 | 0.029 |
| Adjusted $R^2$ | 0.033 | 0.036 | 0.025 | 0.025 | 0.029 |
| Residual Std. Error (df = 71186) | 2.191 | 2.188 | 2.201 | 2.201 | 2.197 |
| F Statistic (df = 8; 71186) | 307.392*** | 336.631*** | 228.676*** | 232.136*** | 262.571*** |
| *Note:* | | | | | *p<0.1; **p<0.05; ***p<0.01 |

**Table S20: Regressing number of children ever born on scores (UKB): matched N and including sex and age controls** The statistical significance does not differ from Table S12

| | *Dependent variable:* | | | | |
|---|---|---|---|---|---|
| | inv_norm_num_children | | | | |
| | (1) | (2) | (3) | (4) | (5) |
| HLMM (matched N) | 0.007** <br> p = 0.028 | | | | |
| Levels mPGS (matched N) | | 0.022*** <br> p = 0.000 | | | |
| Sibling vPGS (matched N) | | | −0.002 <br> p = 0.494 | | |
| Squared-Z (matched N) | | | | 0.002 <br> p = 0.578 | |
| Levene's (matched N) | | | | | −0.002 <br> p = 0.648 |
| PC1 | 0.0005 <br> p = 0.888 | 0.001 <br> p = 0.856 | 0.0004 <br> p = 0.898 | 0.0005 <br> p = 0.897 | 0.0004 <br> p = 0.911 |
| PC2 | 0.001 <br> p = 0.699 | 0.001 <br> p = 0.714 | 0.001 <br> p = 0.692 | 0.001 <br> p = 0.696 | 0.001 <br> p = 0.702 |
| PC3 | −0.005 <br> p = 0.112 | −0.005 <br> p = 0.108 | −0.005 <br> p = 0.110 | −0.005 <br> p = 0.110 | −0.005 <br> p = 0.109 |
| PC4 | 0.002 <br> p = 0.702 | 0.002 <br> p = 0.732 | 0.002 <br> p = 0.704 | 0.002 <br> p = 0.715 | 0.002 <br> p = 0.698 |
| PC5 | 0.012*** <br> p = 0.008 | 0.012*** <br> p = 0.009 | 0.012*** <br> p = 0.006 | 0.012*** <br> p = 0.008 | 0.012*** <br> p = 0.007 |
| Age | 0.021*** <br> p = 0.000 | 0.021*** <br> p = 0.000 | 0.021*** <br> p = 0.000 | 0.021*** <br> p = 0.000 | 0.021*** <br> p = 0.000 |
| Sex | −0.045*** <br> p = 0.000 | −0.045*** <br> p = 0.000 | −0.045*** <br> p = 0.000 | −0.045*** <br> p = 0.000 | −0.045*** <br> p = 0.000 |
| Constant | −0.621*** <br> p = 0.000 | −0.621*** <br> p = 0.000 | −0.621*** <br> p = 0.000 | −0.621*** <br> p = 0.000 | −0.621*** <br> p = 0.000 |
| Observations | 81,282 | 81,282 | 81,282 | 81,282 | 81,282 |
| $R^2$ | 0.031 | 0.032 | 0.031 | 0.031 | 0.031 |
| Adjusted $R^2$ | 0.031 | 0.032 | 0.031 | 0.031 | 0.031 |
| Residual Std. Error (df = 81273) | 0.935 | 0.934 | 0.935 | 0.935 | 0.935 |
| F Statistic (df = 8; 81273) | 329.835*** | 334.778*** | 329.272*** | 329.251*** | 329.238*** |
| *Note:* | | | | | *p<0.1; **p<0.05; ***p<0.01 |

#### S.5 Genetic correlation and heritability analysis

Figures S17 and S18 show a visual representation of the genetic correlation results highlighted in main text Section 2.

**Fig. S17: Bivariate genetic correlation: levels PGS and Squared Z vPGS** The $x$ axis shows the weights for levels of an outcome (mPGS weights). The $y$ axis shows the weights from the Squared Z score method. The correlation close to 1 on the diagonal shows that there is a high degree of overlap in the weights estimated using each method

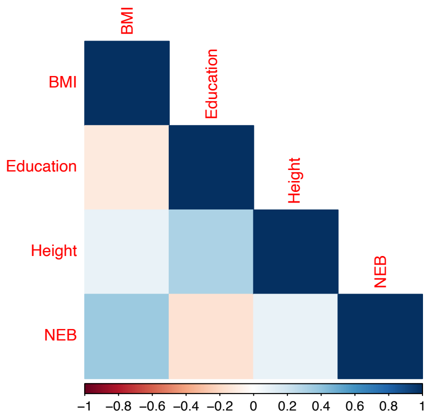


**Fig. S18: Bivariate genetic correlation: levels PGS and vPGS: between outcomes**. The *left panel* shows the bivariate genetic correlation between levels of each outcome variable. The *right panel* shows the bivariate genetic correlation between variability in that outcome, measured using the Squared Z-score vPGS. The two show nearly identical patterns, except for the relationship between BMI and height. While there is a negative genetic correlation between *levels* of BMI and height, there is a positive genetic correlation between *variability in* BMI and height

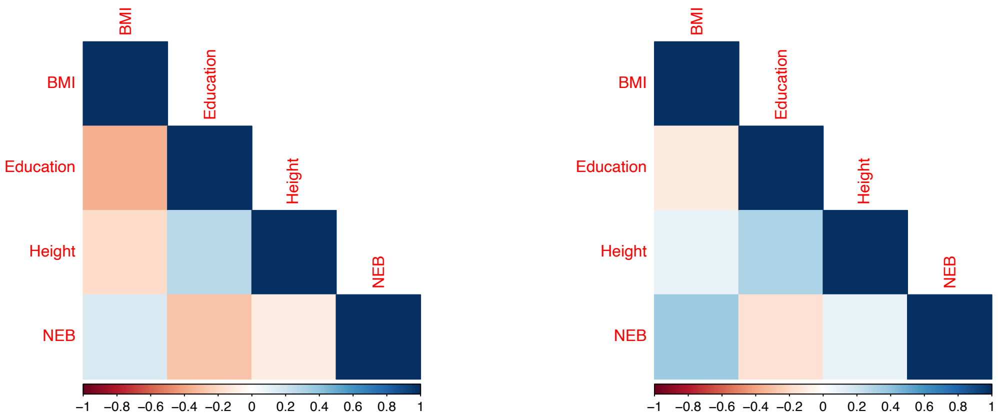


Table S21 shows the analysis of heritability, which are estimated using the better-powered weights: the weights from the full sample for the squared Z-score, HLMM, and Levene's test methods; the weights from the sibling subsample for the sibling SD method. For the sibling SD method, the low heritability could thus result from the smaller sample size.

The first column, in line with past findings, shows the highest heritability for height followed by BMI, education, and finally number of children ever born. We see that plasticity as defined using the Squared Z score and Levene's test have higher heritability, which Section 3.1 shows could be related to capturing levels-relevant genetic contributions; both the sibling standard deviation method and dispersion effects from HLMM capture plasticity in a way that leads to estimates of zero heritability. Interestingly, focusing on the Squared Z estimates, we see that the rank order of heritability changes. That is, levels of height are the most heritable, followed by BMI, then education, and fertility is the least heritable. With the Squared Z method, we find *plasticity in BMI* to be more heritable than plasticity in height.

We also see that LD score regression, for most of the vPGS, does not generate valid estimates of heritability, possibly due to significantly less precise weights than with levels of a trait. The only consistently valid heritability estimates are for the Squared Z score, which we show is highly correlated with the levels PGS. Therefore, we recommend future research focused on developing methods better suited to estimating heritability from variance-focused weights.

**Table S21: Heritability of each outcome: levels versus plasticity** The cells with zero are heritabilities where, due to sampling variability, the heritability was low enough to be close to zero or negative. The estimates are in the cells with the SEs in parantheses.

| | | | Heritability | | |
|---|---|---|---|---|---|
| **Outcome** | Levels | Squared Z | Levene's | Sibling | HLMM |
| Height | 0.40 (0.027) | 0.0028 (0.0024) | Undefined | 0.00 (0.0025) | 0.00 (0.0029) |
| BMI | 0.20 (0.018) | 0.0406 (0.006) | 0.015 (0.004) | 0.00 (0.0024) | 0.00 (0.002) |
| Education | 0.12 (0.005) | 0.0095 (0.003) | 0.28 (0.03) | 0.003 (0.0027) | 0.00 (0.003) |
| Number ever born | 0.03 (0.003) | 0.0033 (0.0028) | Undefined | Undefined | 0.00 (0.0026) |

#### S.6 Details of vPGS validation exercise

In this section, we discuss details of the vPGS validation exercise discussed in main text Section 3.3. First, for examining **within-person BMI variability** within the HRS, we require multiple measurements of BMI.

Figure S19 shows a histogram where the x axis represents the count of BMI observations across the HRS waves and the y axis represents the count of respondents. We see that requiring three or more waves strikes a good balance between (1) retaining the majority of respondents (97% of the total sample) and (2) having a sufficiently high per-respondent set of measurements.

Table S22 shows the results from the analyses of within-person variability, focusing on the raw BMI outcomes. While the dispersion score is the only one that predicts within-person variability at the $p < 0.1$ level, the other scores show trends in the same direction but are perhaps underpowered.

Table S23 shows the results from the analyses of between-person variability operationalized as the squared residual in BMI after regressing levels of BMI on age, sex, and the first 5 PCs. Interesting, the squared Z score, which is specifically designed around this form of variability, is the only one to predict this form of variability.

**Table S22: Relationship between vPGS and within-person variability in BMI in the HRS**

| | *Dependent variable: within-person standard deviation in BMI* | | | |
|---|---|---|---|---|
| | (1) | (2) | (3) | (4) |
| HLMM (matched N) | 0.024 | | | |
| | (0.014) | | | |
| | p = 0.100* | | | |
| Sibling SD (matched N) | | 0.023 | | |
| | | (0.014) | | |
| | | p = 0.109 | | |
| Levene's (matched N) | | | 0.016 | |
| | | | (0.014) | |
| | | | p = 0.280 | |
| Squared Z (matched N) | | | | 0.021 |
| | | | | (0.014) |
| | | | | p = 0.149 |
| Mean Levels | 0.091 | 0.092 | 0.091 | 0.091 |
| | (0.003) | (0.003) | (0.003) | (0.003) |
| | p = 0.000*** | p = 0.000*** | p = 0.000*** | p = 0.000*** |
| Constant | −0.935 | −0.937 | −0.935 | −0.933 |
| | (0.074) | (0.074) | (0.074) | (0.074) |
| | p = 0.000*** | p = 0.000*** | p = 0.000*** | p = 0.000*** |
| Observations | 5,581 | 5,581 | 5,581 | 5,581 |
| $R^2$ | 0.181 | 0.181 | 0.181 | 0.181 |
| Adjusted $R^2$ | 0.181 | 0.181 | 0.180 | 0.181 |
| Residual Std. Error (df = 5578) | 1.074 | 1.074 | 1.074 | 1.074 |
| F Statistic (df = 2; 5578) | 615.955*** | 615.870*** | 615.009*** | 615.572*** |
| *Note:* | | | | *p<0.1; **p<0.05; ***p<0.01 |

**Fig. S19: Number of observations of BMI for each HRS respondent in genetic analytic sample**

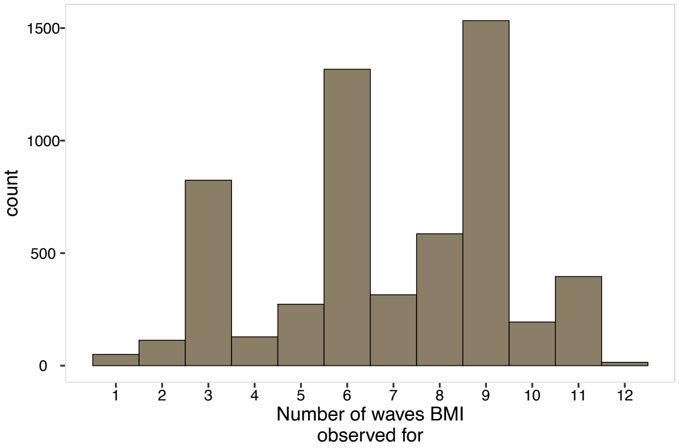


**Table S23: Relationship between vPGS population-level variability in form of squared residual**

| | *Dependent variable: squared residual* | | | |
|---|---|---|---|---|
| | (1) | (2) | (3) | (4) |
| HLMM (matched N) | 0.023<br>(0.023) | | | |
| Sibling SD (matched N) | | 0.017<br>(0.023) | | |
| Squared Z (matched N) | | | 0.047**<br>(0.023) | |
| Levene's (matched N) | | | | 0.028<br>(0.023) |
| Constant | 0.888***<br>(0.023) | 0.888***<br>(0.023) | 0.888***<br>(0.023) | 0.888***<br>(0.023) |
| Observations | 5,744 | 5,744 | 5,744 | 5,744 |
| $R^2$ | 0.0002 | 0.0001 | 0.001 | 0.0003 |
| Adjusted $R^2$ | −0.00001 | −0.0001 | 0.001 | 0.0001 |
| Residual Std. Error (df = 5742) | 1.751 | 1.751 | 1.750 | 1.751 |
| F Statistic (df = 1; 5742) | 0.966 | 0.522 | 4.120** | 1.454 |
| *Note:* | | *p<0.1; **p<0.05; ***p<0.01 | | |

#### S.7 Overlap in top hits

All top hits use a p value threshold of $p < 5 * 10^{-8}$ and examine the top hits from the analysis that matches sample size of the methods that use unrelated individuals to the sample size in the siblings analysis.

**Table S24:** Height: overlap in top hits

| | phenotype | score | ntop_levels | ntop_other | intersect_top | pvals_levels_ifnointersect |
|---|---|---|---|---|---|---|
| 1 | height | hlmm | 64 | 1.00 | 0 | 0.07445 |
| 2 | height | vqtl | 64 | 0.00 | 1 | |
| 3 | height | sib | 64 | 0.00 | 1 | |
| 4 | height | z | 64 | 35.00 | 0 | 0.001889; 0.0008513; 2.705e-05; 0.009572; 0.00 |

**Table S25:** BMI: overlap in top hits

| phenotype | score | ntop_levels | ntop_score | intersect_top | pvals_levels_ifnointersect |
|---|---|---|---|---|---|
| bmi | hlmm | 18 | 1.00 | 0 | 0.528 |
| bmi | vqtl | 18 | 0.00 | 0 | |
| bmi | sib | 18 | 0.00 | 0 | |
| bmi | z | 18 | 200.00 | 0 | 0.01454; 0.0003132; 7.673e-05; 0.009155; 0.06242; 0.0011; 0.5531; 0.1009; 0.007466; 0.01226; 0.02027; 0.00569; 0.08449; 0.2388; 0.2428; 0.0009332; 0.004359; 0.5309; 0.06847; 0.0006797; 0.5245; 0.002594; 0.1336; 0.009217; 0.02783; 0.01883; 0.0008239; 0.08945; 0.07439; 0.004205; 0.01604; 0.9077; 0.007663; 0.013; 0.503; 0.001418; 0.02361; 0.007426; 0.05112; 0.004004; 0.05301; 0.0009327; 0.01109; 0.00597; 0.008897; 0.9922; 0.4357; 0.007316; 0.004316; 0.297; 6.73e-05; 0.02453; 0.004856; 0.00268; 0.07611; 0.0002759; 0.8639; 0.001756; 0.0006801; 0.609; 0.9291; 0.01736; 0.01225; 0.002306; 0.01119; 0.146; 0.007504; 0.003091; 0.002314; 0.007507; 0.0009428; 0.2743; 0.001702; 0.5219; 0.9282; 0.1109; 0.000679; 0.06714; 0.00309; 0.007469; 0.2036; 0.01164; 0.007482; 0.2071; 0.00317; 0.02129; 0.002163; 0.01087; 0.004255; 0.002914; 0.3391; 0.01734; 0.0134; 0.4971; 0.005878; 0.006019; 0.002198; 0.007438; 0.02587; 0.02593; 0.1702; 0.01001; 0.004236; 0.524; 0.008335; 0.1276; 0.4704; 0.7593; 0.002279; 0.6486; 0.002653; 0.007453; 0.03385; 0.01625; 0.0006809; 0.007811; 0.002644; 0.00093; 0.02406; 0.07593; 0.8315; 0.0006708; 0.01242; 0.06782; 0.06526; 0.01765; 0.00747; 0.005772; 0.08778; 0.4903; 0.1493; 0.4002; 0.002827; 0.0006948; 0.004029; 0.004011; 0.2891; 0.0001832; 0.5043; 0.009858; 0.004766; 0.497; 0.004006; 0.007549; 0.002254; 0.1228; 0.07472; 0.05601; 0.007816; 0.208; 0.0004684; 0.005995; 0.5768; 0.02183; 0.2763; 0.3243; 0.04282; 0.01652; 0.007231; 0.09799; 0.003749; 0.2666; 0.01873; 0.0346; 0.7159; 0.001395; 0.0005562; 0.0006783; 0.6943; 0.8948; 0.02638; 0.005871; 0.005215; 0.01738; 0.007478; 0.02597; 0.03142; 0.00402; 0.009968; 0.217; 0.007512; 0.0007039; 0.1933; 0.007471; 0.1465; 0.2322; 0.2568; 0.003403; 0.007717; 0.163; 0.000677; 0.000929; 0.0006802; 0.8408; 0.04026; 0.007501; 0.198; 0.05019; 0.001725; 0.00743 |

**Table S26:** Education: overlap in top hits

| phenotype | score | ntop_levels | ntop_score | intersect_top | pvals_levels_ifnointersect |
|---|---|---|---|---|---|
| education | hlmm | 0 | 3.00 | 0 | 0.529; 0.7218; 0.5367 |
| education | vqtl | 0 | 16574.00 | 0 | Too many to list |
| education | sib | 0 | 0.00 | 0 | |
| education | z | 0 | 0.00 | 0 | |

**Table S27:** Number ever born: overlap in top hits

| phenotype | score | ntop_levels | ntop_score | intersect_top | pvals_levels_ifnointersect |
|---|---|---|---|---|---|
| num_children | hlmm | 0 | 1.00 | 0 | 0.5505 |
| num_children | vqtl | 0 | 23.00 | 0 | 0.002342; 0.4241; 0.2505; 0.05852; 0.0001888; 0.9513; 0.002398; 0.01106; 0.06993; 0.01771; 0.07742; 0.7237; 0.0008457; 0.0004474; 0.02022; 0.04751; 0.002636; 0.9888; 0.2295; 0.08692; 0.002601; 0.002731 |
| num_children | sib | 0 | 0.00 | 0 | |
| num_children | z | 0 | 167.00 | 0 | 0.0099; 0.0794; 0.01299; 0.003631; 0.006495; 0.3128; 0.03401; 0.4733; 0.02076; 0.003443; 0.1982; 0.9477; 0.5937; 0.001317; 0.006689; 0.002595; 0.01875; 0.3659; 0.002411; 0.001093; 0.01114; 0.02072; 0.004717; 0.01082; 0.008148; 0.004973; 0.2402; 0.002586; 0.2679; 0.006315; 0.7825; 0.01467; 0.01042; 0.3483; 0.002431; 0.9415; 0.5721; 0.005348; 0.005211; 0.00953; 0.01806; 0.7311; 0.22; 0.009574; 0.213; 0.9438; 0.5928; 0.3145; 0.4313; 0.2111; 0.3872; 0.004684; 0.008156; 0.2722; 0.1425; 0.4367; 0.02667; 0.009157; 0.05667; 0.01262; 0.9511; 0.5983; 0.5955; 0.0251; 0.6378; 0.4911; 0.003203; 0.5737; 0.963; 0.5146; 0.006448; 0.1042; 0.008184; 0.9246; 0.0005183; 0.4089; 0.01809; 0.1794; 0.1562; 0.1221; 0.3085; 0.002602; 0.01481; 0.0005153; 0.002104; 0.01059; 0.001003; 0.251; 0.004046; 0.6738; 0.7921; 0.004831; 0.003261; 0.794; 0.03536; 0.06875; 0.01837; 0.146; 0.3976; 0.08583; 0.003498; 0.2479; 0.00504; 0.01473; 0.007446; 0.006474; 0.0185; 0.1106; 0.03954; 0.03913; 0.4758; 0.01727; 0.007682; 0.0413; 0.0009807; 0.08617; 0.008771; 0.8459; 0.3327; 0.5986; 0.5383; 0.7082; 0.04961; 0.2861; 0.1987; 0.01327; 0.8427; 0.01806; 0.2237; 0.6982; 0.04345; 0.6464; 0.008871; 0.008838; 0.008779; 0.008722; 0.08277; 0.9129; 0.3834; 0.5638; 0.02626; 0.003635; 0.2126; 0.01486; 0.00854; 0.008702; 0.1376; 0.548; 0.05955; 0.009657; 0.5805; 0.1813; 0.02176; 0.9628; 0.1255; 0.0633; 0.8919; 0.03023; 0.6438; 0.007727; 0.8902; 0.8658; 0.451; 0.09646; 0.8326; 0.008673; 0.04408 |

#### S.8 Additional results: G x E study of heterogeneous impacts of education reform

**Table S28:** Squared Z-Score vPGS, Body Size

| | Continuous | | | Above Threshold | | |
|---|---|---|---|---|---|---|
| | Reduced | 2SLS | | Reduced | 2SLS | |
| | Body Size | Educ16 | Body Size | Body Size | Educ16 | Body Size |
| vPGS | $-0.096^{***}$ | | $-0.106^{\dagger}$ | $-0.024^{***}$ | | $-0.044^{*}$ |
| | (0.007) | | (0.057) | (0.003) | | (0.021) |
| Post Reform | $-0.024^{\dagger}$ | $0.101^{***}$ | | $-0.012^{*}$ | $0.101^{***}$ | |
| | (0.013) | (0.006) | | (0.005) | (0.006) | |
| Educ16 (Instr.) | | | $-0.228^{\dagger}$ | | | $-0.110^{*}$ |
| | | | (0.133) | | | (0.050) |
| mPGS | $0.227^{***}$ | | $0.233^{***}$ | $0.063^{***}$ | | $0.135^{***}$ |
| | (0.007) | | (0.057) | (0.003) | | (0.022) |
| vPGS x Post Reform | −0.001 | | | 0.003 | | |
| | (0.010) | | | (0.004) | | |
| mPGS x Post Reform | −0.004 | | | $-0.015^{***}$ | | |
| | (0.010) | | | (0.004) | | |
| vPGS x Educ16 (Instr.) | | | 0.010 | | | 0.024 |
| | | | (0.065) | | | (0.024) |
| mPGS x Educ16 (Instr.) | | | −0.009 | | | $-0.090^{***}$ |
| | | | (0.065) | | | (0.025) |
| Constant | $-1.885^{***}$ | $0.334^{***}$ | $-1.744^{***}$ | $-0.408^{***}$ | $0.334^{***}$ | $-0.345^{***}$ |
| | (0.256) | (0.101) | (0.263) | (0.097) | (0.101) | (0.100) |
| Observations | 46,736 | 46,719 | 45,961 | 46,736 | 46,719 | 45,961 |
| $R^2$ | 0.061 | 0.066 | 0.061 | 0.025 | 0.066 | 0.025 |
| Adjusted $R^2$ | 0.060 | 0.066 | 0.060 | 0.025 | 0.066 | 0.025 |

*Note:* $\dagger p < 0.1; *p < 0.05; **p < 0.01; ***p < 0.001$

**Table S29:** Levene's vPGS, Body Size

| | Continuous | | | Above Threshold | | |
|---|---|---|---|---|---|---|
| | Reduced | 2SLS | | Reduced | 2SLS | |
| | Body Size | Educ16 | Body Size | Body Size | Educ16 | Body Size |
| vPGS | $-0.082^{***}$ | | $-0.077$ | $-0.019^{***}$ | | $-0.025$ |
| | (0.006) | | (0.049) | (0.002) | | (0.018) |
| Post Reform | $-0.024^{\dagger}$ | $0.101^{***}$ | | $-0.012^{*}$ | $0.101^{***}$ | |
| | (0.013) | (0.006) | | (0.005) | (0.006) | |
| Educ16 (Instr.) | | | $-0.229^{\dagger}$ | | | $-0.111^{*}$ |
| | | | (0.133) | | | (0.050) |
| mPGS | $0.201^{***}$ | | $0.194^{***}$ | $0.056^{***}$ | | $0.117^{***}$ |
| | (0.006) | | (0.049) | (0.002) | | (0.019) |
| vPGS x Post Reform | $-0.006$ | | | $-0.001$ | | |
| | (0.009) | | | (0.003) | | |
| mPGS x Post Reform | $-0.002$ | | | $-0.013^{***}$ | | |
| | (0.009) | | | (0.003) | | |
| vPGS x Educ16 (Instr.) | | | $-0.009$ | | | 0.006 |
| | | | (0.055) | | | (0.021) |
| mPGS x Educ16 (Instr.) | | | 0.006 | | | $-0.077^{***}$ |
| | | | (0.056) | | | (0.021) |
| Constant | $-1.872^{***}$ | $0.334^{***}$ | $-1.729^{***}$ | $-0.405^{***}$ | $0.334^{***}$ | $-0.342^{***}$ |
| | (0.256) | (0.101) | (0.263) | (0.097) | (0.101) | (0.100) |
| Observations | 46,736 | 46,719 | 45,961 | 46,736 | 46,719 | 45,961 |
| $R^2$ | 0.061 | 0.066 | 0.061 | 0.025 | 0.066 | 0.025 |
| Adjusted $R^2$ | 0.060 | 0.066 | 0.060 | 0.025 | 0.066 | 0.025 |

*Note:* $\dagger p < 0.1; *p < 0.05; **p < 0.01; ***p < 0.001$

**Table S30:** HLMM vPGS, Body Size

| | Continuous | | | Above Threshold | | |
|---|---|---|---|---|---|---|
| | Reduced | 2SLS | | Reduced | 2SLS | |
| | Body Size | Educ16 | Body Size | Body Size | Educ16 | Body Size |
| vPGS | $-0.085^{***}$ | | $-0.073$ | $-0.022^{***}$ | | $-0.044^{*}$ |
| | (0.006) | | (0.049) | (0.002) | | (0.018) |
| Post Reform | $-0.023^{\dagger}$ | $0.101^{***}$ | | $-0.012^{*}$ | $0.101^{***}$ | |
| | (0.013) | (0.006) | | (0.005) | (0.006) | |
| Educ16 (Instr.) | | | $-0.225^{\dagger}$ | | | $-0.110^{*}$ |
| | | | (0.133) | | | (0.050) |
| mPGS | $0.202^{***}$ | | $0.198^{***}$ | $0.057^{***}$ | | $0.126^{***}$ |
| | (0.006) | | (0.049) | (0.002) | | (0.019) |
| vPGS x Post Reform | $-0.005$ | | | 0.003 | | |
| | (0.009) | | | (0.003) | | |
| mPGS x Post Reform | $-0.003$ | | | $-0.015^{***}$ | | |
| | (0.009) | | | (0.003) | | |
| vPGS x Educ16 (Instr.) | | | $-0.017$ | | | 0.027 |
| | | | (0.055) | | | (0.021) |
| mPGS x Educ16 (Instr.) | | | 0.003 | | | $-0.087^{***}$ |
| | | | (0.056) | | | (0.021) |
| Constant | $-1.902^{***}$ | $0.334^{***}$ | $-1.764^{***}$ | $-0.410^{***}$ | $0.334^{***}$ | $-0.348^{***}$ |
| | (0.256) | (0.101) | (0.263) | (0.097) | (0.101) | (0.100) |
| Observations | 46,736 | 46,719 | 45,961 | 46,736 | 46,719 | 45,961 |
| $R^2$ | 0.062 | 0.066 | 0.061 | 0.026 | 0.066 | 0.026 |
| Adjusted $R^2$ | 0.061 | 0.066 | 0.061 | 0.025 | 0.066 | 0.025 |

*Note:* $\dagger p < 0.1; *p < 0.05; **p < 0.01; ***p < 0.001$

**Table S31:** Sibling vPGS, Body Size

| | Continuous | | | Above Threshold | | |
|---|---|---|---|---|---|---|
| | Reduced | 2SLS | | Reduced | 2SLS | |
| | Body Size | Educ16 | Body Size | Body Size | Educ16 | Body Size |
| vPGS | $-0.011^{*}$ | | $-0.055$ | $0.005^{*}$ | | 0.011 |
| | (0.005) | | (0.045) | (0.002) | | (0.017) |
| Post Reform | $-0.024^{\dagger}$ | $0.101^{***}$ | | $-0.012^{*}$ | $0.101^{***}$ | |
| | (0.013) | (0.006) | | (0.005) | (0.006) | |
| Educ16 (Instr.) | | | $-0.233^{\dagger}$ | | | $-0.112^{*}$ |
| | | | (0.134) | | | (0.051) |
| mPGS | $0.168^{***}$ | | $0.166^{***}$ | $0.047^{***}$ | | $0.105^{***}$ |
| | (0.005) | | (0.045) | (0.002) | | (0.017) |
| vPGS x Post Reform | 0.012 | | | 0.0004 | | |
| | (0.008) | | | (0.003) | | |
| mPGS x Post Reform | $-0.005$ | | | $-0.013^{***}$ | | |
| | (0.008) | | | (0.003) | | |
| vPGS x Educ16 (Instr.) | | | 0.057 | | | $-0.007$ |
| | | | (0.052) | | | (0.020) |
| mPGS x Educ16 (Instr.) | | | $-0.001$ | | | $-0.073^{***}$ |
| | | | (0.051) | | | (0.019) |
| Constant | $-1.871^{***}$ | $0.334^{***}$ | $-1.731^{***}$ | $-0.402^{***}$ | $0.334^{***}$ | $-0.339^{***}$ |
| | (0.257) | (0.101) | (0.264) | (0.097) | (0.101) | (0.100) |
| Observations | 46,736 | 46,719 | 45,961 | 46,736 | 46,719 | 45,961 |
| $R^2$ | 0.053 | 0.066 | 0.053 | 0.023 | 0.066 | 0.022 |
| Adjusted $R^2$ | 0.052 | 0.066 | 0.052 | 0.022 | 0.066 | 0.022 |

*Note:* $\dagger p < 0.1; *p < 0.05; **p < 0.01; ***p < 0.001$

**Table S32:** Squared Z-Score vPGS, Body Size, No mPGS

| | Continuous | | | Above Threshold | | |
|---|---|---|---|---|---|---|
| | Reduced | 2SLS | | Reduced | 2SLS | |
| | Body Size | Educ16 | Body Size | Body Size | Educ16 | Body Size |
| vPGS | 0.047*** | | 0.057 | 0.015*** | | 0.044** |
| | (0.005) | | (0.045) | (0.002) | | (0.017) |
| Post Reform | −0.024† | 0.101*** | | −0.012* | 0.101*** | |
| | (0.014) | (0.006) | | (0.005) | (0.006) | |
| Educ16 (Instr.) | | | −0.235† | | | −0.112* |
| | | | (0.136) | | | (0.051) |
| vPGS x Post Reform | −0.006 | | | −0.007* | | |
| | (0.008) | | | (0.003) | | |
| vPGS x Educ16 (Instr.) | | | −0.016 | | | −0.037† |
| | | | (0.052) | | | (0.019) |
| Constant | −1.811*** | 0.334*** | −1.665*** | −0.391*** | 0.334*** | −0.327** |
| | (0.262) | (0.101) | (0.269) | (0.098) | (0.101) | (0.101) |
| Observations | 46,736 | 46,719 | 45,961 | 46,736 | 46,719 | 45,961 |
| $R^2$ | 0.019 | 0.066 | 0.019 | 0.007 | 0.066 | 0.006 |
| Adjusted $R^2$ | 0.018 | 0.066 | 0.018 | 0.006 | 0.066 | 0.006 |

*Note:* $\dagger p < 0.1; *p < 0.05; **p < 0.01; ***p < 0.001$

**Table S33:** Levene's vPGS, Body Size, No mPGS

| | Continuous | | | Above Threshold | | |
|---|---|---|---|---|---|---|
| | Reduced | 2SLS | | Reduced | 2SLS | |
| | Body Size | Educ16 | Body Size | Body Size | Educ16 | Body Size |
| vPGS | 0.003 | | 0.020 | $0.004^{*}$ | | 0.027 |
| | (0.005) | | (0.045) | (0.002) | | (0.017) |
| Post Reform | $-0.024^{\dagger}$ | $0.101^{***}$ | | $-0.012^{*}$ | $0.101^{***}$ | |
| | (0.014) | (0.006) | | (0.005) | (0.006) | |
| Educ16 (Instr.) | | | $-0.232^{\dagger}$ | | | $-0.111^{*}$ |
| | | | (0.136) | | | (0.051) |
| vPGS x Post Reform | $-0.008$ | | | $-0.007^{*}$ | | |
| | (0.008) | | | (0.003) | | |
| vPGS x Educ16 (Instr.) | | | $-0.024$ | | | $-0.030$ |
| | | | (0.052) | | | (0.019) |
| Constant | $-1.809^{***}$ | $0.334^{***}$ | $-1.662^{***}$ | $-0.391^{***}$ | $0.334^{***}$ | $-0.326^{**}$ |
| | (0.262) | (0.101) | (0.269) | (0.098) | (0.101) | (0.101) |
| Observations | 46,736 | 46,719 | 45,961 | 46,736 | 46,719 | 45,961 |
| $R^2$ | 0.016 | 0.066 | 0.016 | 0.005 | 0.066 | 0.005 |
| Adjusted $R^2$ | 0.016 | 0.066 | 0.016 | 0.004 | 0.066 | 0.004 |

*Note:* $\dagger p < 0.1; *p < 0.05; **p < 0.01; ***p < 0.001$

**Table S34:** HLMM vPGS, Body Size, No mPGS

| | Continuous | | | Above Threshold | | |
|---|---|---|---|---|---|---|
| | Reduced | 2SLS | | Reduced | 2SLS | |
| | Body Size | Educ16 | Body Size | Body Size | Educ16 | Body Size |
| vPGS | −0.001 | | 0.028 | 0.002 | | 0.013 |
| | (0.005) | | (0.045) | (0.002) | | (0.017) |
| Post Reform | $-0.024^{\dagger}$ | $0.101^{***}$ | | $-0.012^{*}$ | $0.101^{***}$ | |
| | (0.014) | (0.006) | | (0.005) | (0.006) | |
| Educ16 (Instr.) | | | $-0.232^{\dagger}$ | | | $-0.111^{*}$ |
| | | | (0.136) | | | (0.051) |
| vPGS x Post Reform | −0.009 | | | −0.003 | | |
| | (0.008) | | | (0.003) | | |
| vPGS x Educ16 (Instr.) | | | −0.038 | | | −0.014 |
| | | | (0.052) | | | (0.019) |
| Constant | $-1.812^{***}$ | $0.334^{***}$ | $-1.665^{***}$ | $-0.390^{***}$ | $0.334^{***}$ | $-0.326^{**}$ |
| | (0.262) | (0.101) | (0.269) | (0.098) | (0.101) | (0.101) |
| Observations | 46,736 | 46,719 | 45,961 | 46,736 | 46,719 | 45,961 |
| $R^2$ | 0.016 | 0.066 | 0.016 | 0.005 | 0.066 | 0.005 |
| Adjusted $R^2$ | 0.016 | 0.066 | 0.016 | 0.004 | 0.066 | 0.004 |

*Note:* $\dagger p < 0.1; *p < 0.05; **p < 0.01; ***p < 0.001$

**Table S35:** Sibling vPGS, Body Size, No mPGS

| | Continuous | | | Above Threshold | | |
|---|---|---|---|---|---|---|
| | Reduced | 2SLS | | Reduced | 2SLS | |
| | Body Size | Educ16 | Body Size | Body Size | Educ16 | Body Size |
| vPGS | 0.008 | | −0.048 | 0.010*** | | 0.020 |
| | (0.005) | | (0.046) | (0.002) | | (0.017) |
| Post Reform | −0.024† | 0.101*** | | −0.012* | 0.101*** | |
| | (0.014) | (0.006) | | (0.005) | (0.006) | |
| Educ16 (Instr.) | | | −0.232† | | | −0.111* |
| | | | (0.136) | | | (0.051) |
| vPGS x Post Reform | 0.014† | | | −0.0004 | | |
| | (0.008) | | | (0.003) | | |
| vPGS x Educ16 (Instr.) | | | 0.072 | | | −0.012 |
| | | | (0.053) | | | (0.020) |
| Constant | −1.799*** | 0.334*** | −1.653*** | −0.384*** | 0.334*** | −0.320** |
| | (0.262) | (0.101) | (0.269) | (0.098) | (0.101) | (0.101) |
| Observations | 46,736 | 46,719 | 45,961 | 46,736 | 46,719 | 45,961 |
| $R^2$ | 0.017 | 0.066 | 0.017 | 0.006 | 0.066 | 0.006 |
| Adjusted $R^2$ | 0.016 | 0.066 | 0.016 | 0.005 | 0.066 | 0.005 |
| *Note:* | | | | | | $\dagger p < 0.1; *p < 0.05; **p < 0.01; ***p < 0.001$ |

**Table S36: Squared Z-Score vPGS, Education Outcomes**

| | Left School 16 or later | | Certification | | O-levels or CSE | | A-levels | |
|---|---|---|---|---|---|---|---|---|
| Controls? | N | Y | N | Y | N | Y | N | Y |
| | (1) | (2) | (3) | (4) | (5) | (6) | (7) | (8) |
| vPGS | 0.015*** | −0.024*** | 0.008*** | −0.024*** | −0.016*** | 0.004 | 0.001 | −0.005† |
| | (0.002) | (0.002) | (0.002) | (0.002) | (0.003) | (0.003) | (0.002) | (0.002) |
| Post Reform | 0.152*** | 0.119*** | 0.074*** | 0.028*** | 0.087*** | 0.062*** | 0.018*** | 0.007 |
| | (0.003) | (0.005) | (0.003) | (0.005) | (0.004) | (0.007) | (0.003) | (0.005) |
| mPGS | | 0.073*** | | 0.060*** | | −0.036*** | | 0.012*** |
| | | (0.002) | | (0.002) | | (0.003) | | (0.002) |
| vPGS x Post Reform | −0.012*** | 0.020*** | 0.001 | 0.018*** | −0.001 | 0.020*** | −0.001 | 0.001 |
| | (0.003) | (0.003) | (0.003) | (0.003) | (0.004) | (0.005) | (0.003) | (0.004) |
| mPGS x Post Reform | | −0.060*** | | −0.033*** | | −0.040*** | | −0.003 |
| | | (0.003) | | (0.003) | | (0.005) | | (0.004) |
| Constant | 0.819*** | 0.690*** | 0.864*** | 0.265** | 0.278*** | 0.366** | 0.124*** | −0.049 |
| | (0.002) | (0.094) | (0.002) | (0.091) | (0.003) | (0.139) | (0.002) | (0.103) |
| Observations | 46,719 | 46,719 | 47,474 | 47,474 | 47,518 | 47,518 | 47,474 | 47,474 |
| $R^2$ | 0.057 | 0.087 | 0.015 | 0.045 | 0.010 | 0.030 | 0.001 | 0.004 |
| Adjusted $R^2$ | 0.057 | 0.086 | 0.015 | 0.044 | 0.010 | 0.029 | 0.001 | 0.003 |

*Note:* *p<0.1; **p<0.05; ***p<0.01
$\dagger p < 0.1; *p < 0.05; **p < 0.01; ***p < 0.001$

#### Table S37: Levene's vPGS, Education Outcomes

| | Left School 16 or later | | Certification | | O-levels or CSE | | A-levels | |
|---|---|---|---|---|---|---|---|---|
| Controls? | N | Y | N | Y | N | Y | N | Y |
| | (1) | (2) | (3) | (4) | (5) | (6) | (7) | (8) |
| vPGS | 0.028*** | −0.017*** | 0.020*** | −0.016*** | −0.019*** | 0.008* | 0.004† | −0.003 |
| | (0.002) | (0.002) | (0.002) | (0.002) | (0.003) | (0.004) | (0.002) | (0.003) |
| Post Reform | 0.152*** | 0.119*** | 0.074*** | 0.028*** | 0.087*** | 0.062*** | 0.018*** | 0.007 |
| | (0.003) | (0.005) | (0.003) | (0.005) | (0.004) | (0.007) | (0.003) | (0.005) |
| mPGS | | 0.071*** | | 0.057*** | | −0.038*** | | 0.011*** |
| | | (0.002) | | (0.002) | | (0.004) | | (0.003) |
| vPGS x Post Reform | −0.021*** | 0.018*** | −0.007* | 0.013*** | −0.012** | 0.016** | 0.001 | 0.004 |
| | (0.003) | (0.004) | (0.003) | (0.004) | (0.004) | (0.006) | (0.003) | (0.004) |
| mPGS x Post Reform | | −0.060*** | | −0.032*** | | −0.039*** | | −0.005 |
| | | (0.004) | | (0.004) | | (0.006) | | (0.004) |
| Constant | 0.818*** | 0.688*** | 0.864*** | 0.263** | 0.278*** | 0.363** | 0.124*** | −0.050 |
| | (0.002) | (0.094) | (0.002) | (0.091) | (0.003) | (0.139) | (0.002) | (0.103) |
| Observations | 46,719 | 46,719 | 47,474 | 47,474 | 47,518 | 47,518 | 47,474 | 47,474 |
| $R^2$ | 0.061 | 0.086 | 0.018 | 0.043 | 0.011 | 0.029 | 0.001 | 0.004 |
| Adjusted $R^2$ | 0.061 | 0.085 | 0.018 | 0.042 | 0.011 | 0.029 | 0.001 | 0.003 |

*Note:* *p<0.1; **p<0.05; ***p<0.01

$\dagger p < 0.1; *p < 0.05; **p < 0.01; ***p < 0.001$

**Table S38: HLMM vPGS, Education Outcomes**

| | Left School 16 or later | | Certification | | O-levels or CSE | | A-levels | |
|---|---|---|---|---|---|---|---|---|
| Controls? | N | Y | N | Y | N | Y | N | Y |
| | (1) | (2) | (3) | (4) | (5) | (6) | (7) | (8) |
| vPGS | −0.026*** | −0.012*** | −0.030*** | −0.016*** | −0.005 | −0.002 | −0.026*** | −0.014*** |
| | (0.003) | (0.003) | (0.003) | (0.003) | (0.004) | (0.004) | (0.004) | (0.004) |
| Post Reform | 0.156*** | 0.143*** | 0.055*** | 0.053*** | 0.063*** | 0.060*** | −0.008 | −0.007 |
| | (0.008) | (0.008) | (0.008) | (0.008) | (0.011) | (0.011) | (0.009) | (0.009) |
| mPGS | | 0.050*** | | 0.051*** | | 0.004 | | 0.046*** |
| | | (0.003) | | (0.003) | | (0.004) | | (0.004) |
| vPGS x Post Reform | 0.019*** | 0.008† | 0.012* | 0.005 | 0.009 | 0.001 | 0.003 | 0.004 |
| | (0.004) | (0.005) | (0.005) | (0.005) | (0.006) | (0.006) | (0.005) | (0.006) |
| mPGS x Post Reform | | −0.040*** | | −0.027*** | | −0.027*** | | −0.00002 |
| | | (0.005) | | (0.005) | | (0.006) | | (0.006) |
| Constant | 0.774*** | 0.466** | 0.804*** | −0.010 | 0.556*** | 0.257 | 0.248*** | −0.267 |
| | (0.004) | (0.147) | (0.005) | (0.156) | (0.006) | (0.206) | (0.005) | (0.178) |
| Observations | 25,690 | 25,690 | 26,012 | 26,012 | 26,012 | 26,012 | 26,012 | 26,012 |
| $R^2$ | 0.087 | 0.115 | 0.043 | 0.070 | 0.017 | 0.027 | 0.004 | 0.023 |
| Adjusted $R^2$ | 0.087 | 0.114 | 0.043 | 0.069 | 0.017 | 0.025 | 0.004 | 0.021 |

*Note:* $\dagger p < 0.1; *p < 0.05; **p < 0.01; ***p < 0.001$

**Table S39: Sibling vPGS, Education Outcomes**

| | Left School 16 or later | | Certification | | O-levels or CSE | | A-levels | |
|---|---|---|---|---|---|---|---|---|
| Controls? | N | Y | N | Y | N | Y | N | Y |
| | (1) | (2) | (3) | (4) | (5) | (6) | (7) | (8) |
| vPGS | 0.001 | 0.008*** | −0.001 | 0.005** | 0.001 | −0.001 | 0.001 | 0.002 |
| | (0.002) | (0.002) | (0.002) | (0.002) | (0.003) | (0.003) | (0.002) | (0.002) |
| Post Reform | 0.119*** | 0.113*** | 0.028*** | 0.028*** | 0.028*** | 0.028*** | 0.007 | 0.007 |
| | (0.005) | (0.005) | (0.005) | (0.005) | (0.007) | (0.007) | (0.005) | (0.005) |
| mPGS | | 0.061*** | | 0.047*** | | −0.020*** | | 0.009*** |
| | | (0.002) | | (0.002) | | (0.003) | | (0.002) |
| vPGS x Post Reform | −0.002 | −0.008** | −0.002 | −0.004 | 0.0003 | −0.001 | 0.006† | 0.005† |
| | (0.003) | (0.003) | (0.003) | (0.003) | (0.004) | (0.004) | (0.003) | (0.003) |
| mPGS x Post Reform | | −0.050*** | | −0.023*** | | −0.011** | | −0.002 |
| | | (0.003) | | (0.003) | | (0.004) | | (0.003) |
| Constant | 0.836*** | 0.679*** | 0.889*** | 0.246** | 0.222*** | 0.249† | 0.130*** | −0.057 |
| | (0.003) | (0.093) | (0.003) | (0.091) | (0.004) | (0.129) | (0.003) | (0.103) |
| Observations | 46,719 | 46,719 | 47,474 | 47,474 | 47,474 | 47,474 | 47,474 | 47,474 |
| $R^2$ | 0.057 | 0.091 | 0.017 | 0.043 | 0.002 | 0.011 | 0.001 | 0.004 |
| Adjusted $R^2$ | 0.057 | 0.090 | 0.017 | 0.042 | 0.002 | 0.010 | 0.001 | 0.003 |

*Note:* $\dagger p < 0.1; *p < 0.05; **p < 0.01; ***p < 0.001$
